# Comparing Bayesian and non-Bayesian accounts of human confidence reports

**DOI:** 10.1101/093203

**Authors:** William T. Adler, Wei Ji Ma

## Abstract

Humans can meaningfully report their confidence in a perceptual or cognitive decision. It is widely believed that these reports reflect the Bayesian probability that the decision is correct, but this hypothesis has not been rigorously tested against non-Bayesian alternatives. We use two perceptual categorization tasks in which Bayesian confidence reporting requires subjects to take sensory uncertainty into account in a specific way. We find that subjects do take sensory uncertainty into account when reporting confidence, suggesting that brain areas involved in reporting confidence can access low-level representations of sensory uncertainty. However, behavior is not fully consistent with the Bayesian hypothesis and is better described by simple heuristic models. Both conclusions are robust to changes in the uncertainty manipulation, task, response modality, model comparison metric, and additional flexibility in the Bayesian model. Our results suggest that adhering to a rational account of confidence behavior may require incorporating implementational constraints.

## Introduction

People often have a sense of a level of confidence about their decisions. Such a “feeling of knowing”^15,51^ may serve to improve performance in subsequent decisions^61^, learning^51^, and group decision-making^7^. Much recent work has focused on identifying brain regions and neural mechanisms responsible for the computation of confidence in humans^22,23,67^, nonhuman primates^21,37,40^, and rodents^35^. In the search for the neural correlates of confidence, the leading premise has been that confidence is Bayesian, i.e., the observer’s estimated probability that a choice is correct^20,34,51,62^. In human studies, however, naÏve subjects can give a meaningful answer when you ask them to rate their confidence about a decision^59^; thus, “confidence” intrinsically means something to people, and it is not a foregone conclusion that this intrinsic sense corresponds to the Bayesian definition. Therefore, we regard the above “definition” as a testable hypothesis about the way the brain computes explicit confidence reports; we use Bayesian decision theory to formalize this hypothesis.

Bayesian decision theory provides a general and often quantitatively accurate account of perceptual decisions in a wide variety of tasks^39,41,47^. According to this theory, the decision-maker combines knowledge about the statistical structure of the world with the present sensory input to compute a posterior probability distribution over possible states of the world. In principle, a confidence report might be derived from the same posterior distribution; this is the hypothesis described above, which we will call the Bayesian confidence hypothesis (BCH). The main goal of this paper is to test that hypothesis. Recent studies have attempted to test the BCH^5,69^ but, because of their experimental designs, are unable to meaningfully distinguish the Bayesian model from any other model of confidence.

Recent work has proposed possible qualitative signatures of Bayesian confidence^31^. However, the observation (or lack thereof) of these signatures provides an uncertain amount of evidence in favor of (or against) the Bayesian model, and the signatures are therefore not useful for determining which computations underlie confidence reports^4^. To objectively and quantitatively determine whether confidence ratings appear to be Bayesian, we use a formal model comparison approach. We test the predictions of the BCH as we vary the quality of the sensory evidence and the task structure within individuals. We compare Bayesian models against a variety of alternative models, something that is rarely done but very important for the epistemological standing of Bayesian claims^11,32^. We find that the BCH qualitatively describes human behavior but that quantitatively, even the most flexible Bayesian model is outperformed by models that take uncertainty into account in a non-Bayesian way.

## Results

### Experiment 1

During each session, each subject completed two orientation categorization tasks, Tasks A and B. On each trial, a category *C* was selected randomly (both categories were equally probable), and a stimulus *s* was drawn from the corresponding stimulus distribution and displayed. The subject categorized the stimulus and simultaneously reported their confidence on a 4-point scale, with a single button press (Figure 1a). Using a single button press for choice and confidence prevented post-choice influences on the confidence judgment^53^ and emphasized that confidence should reflect the observer’s perception rather than a preceding motor response. The categories were defined by normal distributions on orientation, which differed by task (Figure 1b). In Task A, the distributions had different means (±*μ*_*C*_) and the same standard deviation (*σC*); leftward-tilting stimuli were more likely to be from category 1. Variants of Task A are common in decisionmaking studies^14^. In Task B, the distributions had the same mean (0°) and different standard deviations (*σ* _1_, *σ*2); stimuli around the horizontal were more likely to be from category 1. Variants of Task B are less common^44,63,68^ but have some properties of perceptual organization tasks; for example, a subject may have to detect when a stimulus belongs to a narrow category (e.g., in which two line segments are collinear) that is embedded in a a broader category (e.g., in which two line segments are unrelated).

**Figure 1:**
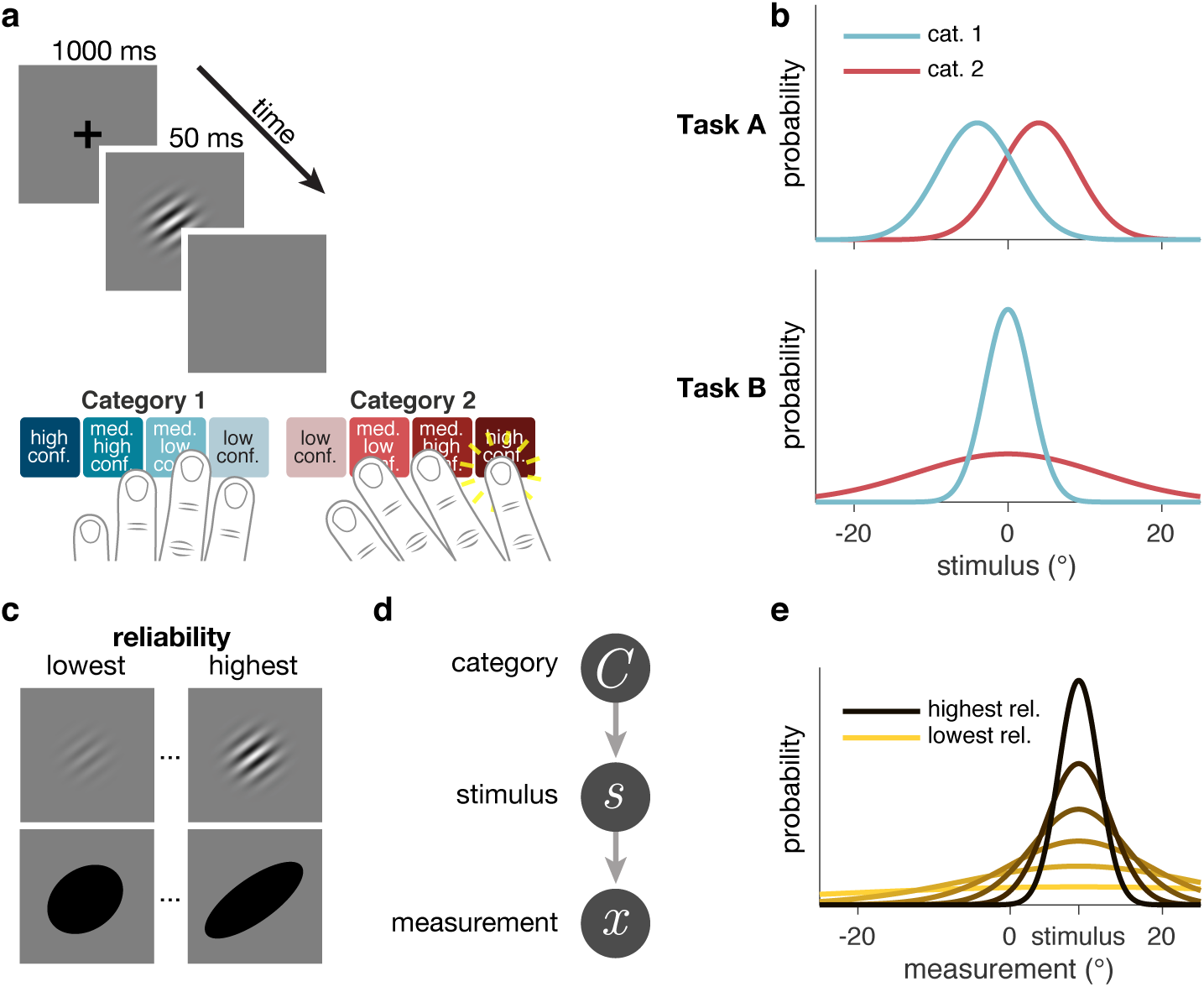
Task design. (**a**) Schematic of a test block trial. After stimulus offset, subjects reported category and confidence level with a single button press. (**b**) Stimulus distributions for Tasks A and B. (**c**) Examples of low and high reliability stimuli. Six (out of eleven) subjects saw drifting Gabors, and five subjects saw ellipses. (**d**) Generative model. (**e**) Example measurement distributions at different reliability levels. In all models (except Linear Neural), the measurement is assumed to be drawn from a Gaussian distribution centered on the true stimulus, with s.d. dependent on reliability.

Subjects were highly trained on the categories; during training, we only used highest-reliability stimuli, and we provided trial-to-trial category correctness feedback. Subjects were then tested with 6 different reliability levels, which were chosen randomly on each trial. During testing, correctness feedback was withheld to avoid the possibility that confidence simply reflects a learned mapping between stimuli and the probability of being correct, something that no other confidence studies have done^42,49,63^.

Because we are interested in subjects’ intrinsic computation of confidence, we did not instruct or incentivize them to assign probability ranges to each button (e.g., by using a scoring rule^13,29,50^). If we had, we would have essentially been training subjects to use a specific model of confidence.

To ensure that our results were independent of stimulus type, we used two kinds of stimuli. Some subjects saw oriented drifting Gabors; for these subjects, stimulus reliability was manipulated through contrast. Other subjects saw oriented ellipses; for these subjects, stimulus reliability was manipulated through ellipse elongation (Figure 1c). We found no major differences in model rankings between Gabor and ellipse subjects, therefore we will make no distinctions between the groups (Supplementary Information).

For modeling purposes, we assume that the observer’s internal representation of the stimulus is a noisy measurement *x*, drawn from a Gaussian distribution with mean *s* and s.d. *σ* (Figure 1d,e). In the model, *σ* (i.e., uncertainty) is a fitted function of stimulus reliability (Supplementary Information).

### Bayesian model

A Bayes-optimal observer uses knowledge of the generative model to make a decision that maximizes the probability of being correct. Here, when the measurement on a given trial is *x*, this strategy amounts to choosing the category *C* for which the posterior probability *p*(*C* | *x*) is highest. This is equivalent to reporting category 1 when the log posterior ratio, 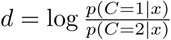, is positive.

In Task A, *d* is 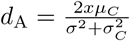. Therefore, the ideal observer reports category 1 when *x* is positive; this is the structure of many psychophysical tasks. In Task *B*, however, *d* is 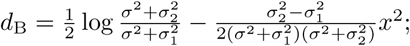 the observer needs both *x* and *σ* in order to make an optimal decision.

From the point of view of the observer, *σ* is the trial-to-trial level of sensory uncertainty associated with the measurement^45^. In a minor variation of the optimal observer, we allow for the possibility that the observer’s prior belief over category, *p*(*C*), is different from the true value of (0.5,0.5); this adds a constant to *d*_A_ and *d*_B_.

We introduce the Bayesian confidence hypothesis (BCH), stating that confidence reports depend on the internal representation of the stimulus (here *x*) only via *d.* In the BCH, the observer chooses a response by comparing *d* to a set of category and confidence boundaries. For example, whenever *d* falls within a certain range, the observer presses the “medium-low confidence, category 2” button. The BCH is thus an extension of the choice model described above, wherein the value of *d* is used to compute confidence as well as chosen category. There is another way of thinking about this. Bayesian models assume that subjects compute *d* in order to make an optimal choice. Assuming people compute *d* at all, are they able to use it to report confidence as well? We refer to the Bayesian model here as simply “Bayes.” We also tested several more constrained versions of this model (Supplementary Information).

The observer’s decision can be summarized as a mapping from a combination of a measurement and an uncertainty level (*x, σ*) to a response that indicates both category and confidence. We can visualize this mapping as in Figure 2, first column. It is clear that the pattern of decision boundaries in the BCH is qualitatively very different between Task A and Task B. In Task A, the decision boundaries are quadratic functions of uncertainty; confidence decreases monotonically with uncertainty and increases with the distance of the measurement from 0. In Task B, the decision boundaries are neither linear nor quadratic.

**Figure 2:**
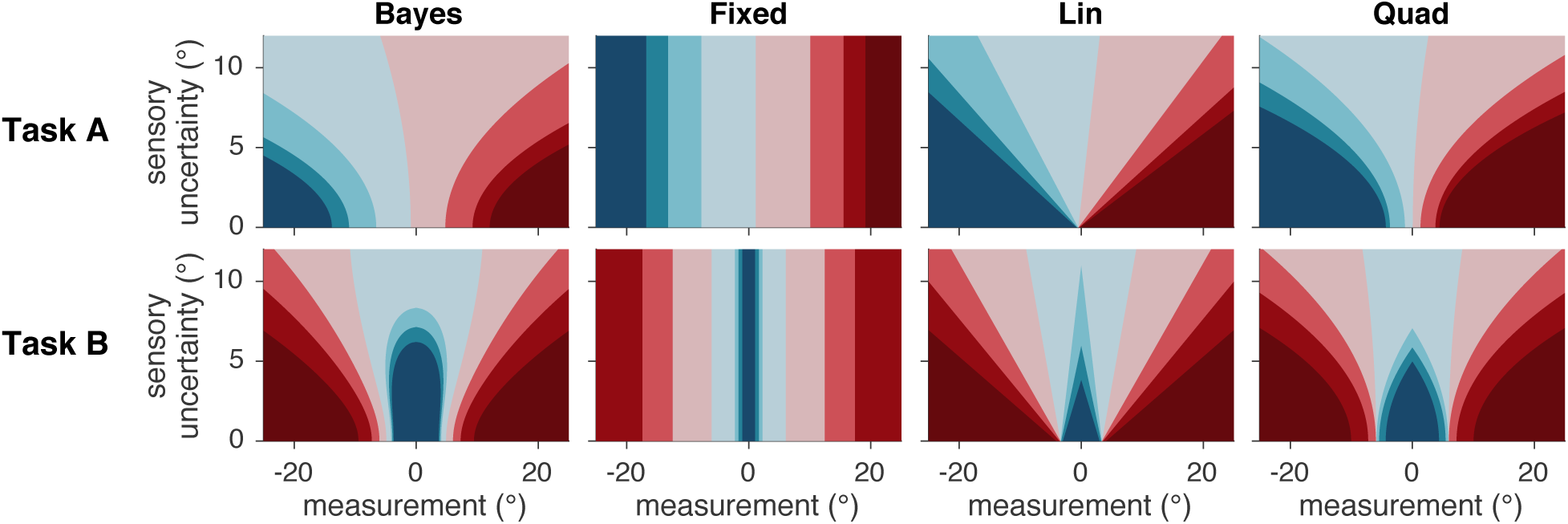
Decision rules/mappings in four models. Each model corresponds to a different mapping from a measurement and uncertainty level to a category and confidence response. Colors correspond to category and confidence response, as in Figure 1a. Plots were generated from the mean of subject 4’s posterior distribution over parameters. These figures should be used only as an aid for understanding the models’ decision rules, not for closely interpreting the different fitted rules across models; interpretation is complicated by, among other considerations, the fact that some regions have very few trials.

### Alternative models

At first glance, it seems obvious that sensory uncertainty is relevant to the computation of confidence. However, this is by no means a given; in fact, a prominent proposal is that confidence is based on the distance between the measurement and the decision boundary, without any role for sensory uncertainty^35,40,64^. Therefore, we tested a model (Fixed) in which the response is a function of the measurement alone (equivalent to a maximum likelihood estimate of the stimulus orientation), and not of the uncertainty of that measurement (Figure 2, second column).

We also tested heuristic models in which the subject uses their knowledge of their sensory uncertainty but does not compute a posterior distribution over category. We have previously classified such models as *probabilistic non-Bayesian*^46^. In the Orientation Estimation model, subjects base their response on a maximum a posteriori estimate of orientation (rather than category), using the mixture of the two stimulus distributions as a prior distribution. In the Linear Neural model, subjects base their response on a linear function of the output of a hypothetical population of neurons.

We derived two additional probabilistic non-Bayesian models, Lin and Quad, from the observation that the Bayesian decision criteria are an approximately linear function of uncertainty in some measurement regimes and approximately quadratic in others. These models are able to produce approximately Bayesian behavior without actually performing any computation of the posterior. In Lin and Quad, subjects base their response on a linear or a quadratic function of *x* and *σ*, respectively (Supplementary Information). A comparison of the Lin and Quad columns to the Bayes column in Figure 2 demonstrates that Lin and Quad can approximate the Bayesian mapping from (*x, σ*) to response despite not being based on the Bayesian decision variable. All of the models we tested were variants of the six models described so far (Bayes, Fixed, Orientation Estimation, Linear Neural, Lin, Quad).

Each trial consists of the experimentally determined orientation and reliability level and the subject’s category and confidence response (an integer between 1 and 8). This is a very rich data set, which we summarize in Figure 3. We find the following effects: performance and confidence increase as a function of reliability (Figure 3a,b,h,i), and high-confidence reports are less frequent than low-confidence reports (Figure 3e,f). Note Figure 3c,d especially; this is the projection of the data that we will use to demonstrate model fits for the rest of this paper. We use this projection because the vertical axis (mean button press) most closely approximates the form of the raw data. Additionally, because our models are differentiated by how they use uncertainty, it is informative to plot how response changes as a function of reliability, in addition to category and task.

**Figure 3:**
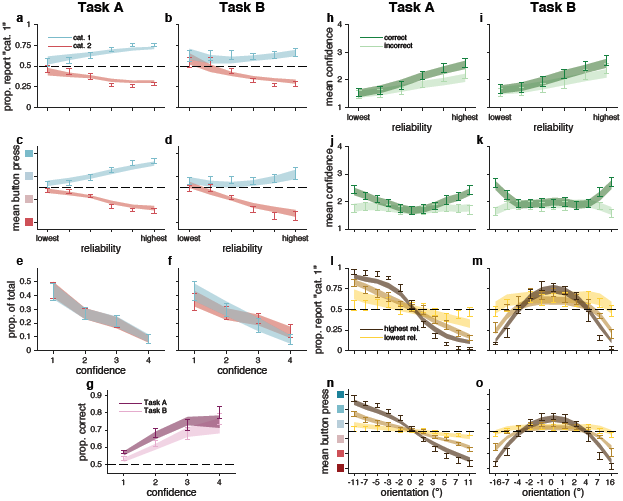
Behavioral data and fits from best model (Quad), experiment 1. Error bars represent ±1 s.e.m. across 11 subjects. Shaded regions represent ±1 s.e.m. on model fits ( (Supplementary Information)). (**a,b**) Proportion “category 1” reports as a function of stimulus reliability and true category. (**c,d**) Mean button press as a function of stimulus reliability and true category. (**e,f**) Normalized histogram of confidence reports for both true categories. (**g**) Proportion correct category reports as a function of confidence report and task. (**h,i**) Mean confidence as a function of stimulus reliability and correctness. (**j,k**) Mean confidence as a function of stimulus orientation and reliability. (**l,m**) Proportion “category 1” reports as a function of stimulus orientation and reliability. (**n,o**) Mean button press as a function of stimulus orientation and reliability. (**c,d,n,o**) Vertical axis label colors correspond to button presses, as in Figure 1a. (**l–o**) For clarity, only 3 of 6 reliability levels are shown, although models were fit to all reliability levels.

### Model comparison

We used Markov Chain Monte Carlo (MCMC) sampling to fit models to raw individual-subject data. To account for overfitting, we compared models using leave-one-out cross-validated log likelihood scores (LOO) computed with the full posteriors obtained through MCMC^72^. A model recovery analysis ensured that our models are meaningfully distinguishable (Figure S5). Unless otherwise noted, models were fit jointly to Task A and B category and confidence responses (Supplementary Information).

#### Use of sensory uncertainty

We first compared Bayes to the Fixed model, in which the observer does not take trial-to-trial sensory uncertainty into account (Figure 4). Fixed provides a poor fit to the data, indicating that observers use not only a point estimate of their measurement, but also their uncertainty about that measurement. Bayes outperforms Fixed by a summed LOO difference (median and 95% CI of bootstrapped sums across subjects) of 2265 [498, 4253]. For the rest of this paper, we will report model comparison results using this format (Section 4.7.2).

**Figure 4:**
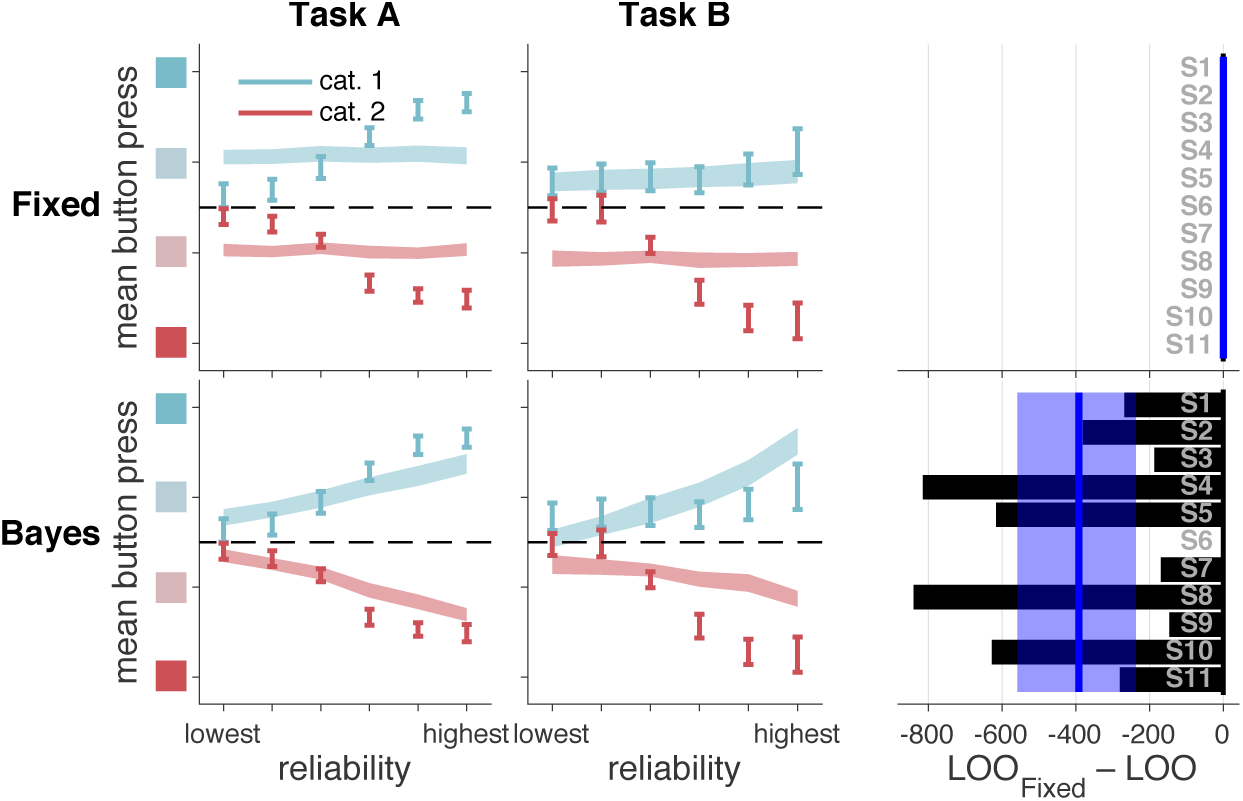
Model fits and model comparison for models Fixed and Bayes. Bayes provides a better fit, but both models have large deviations from the data. Left and middle columns: model fits to mean button press as a function of reliability, true category, and task. Error bars represent ±1 s.e.m. across 11 subjects. Shaded regions represent ±1 s.e.m. on model fits, with each model on a separate row. Right column: LOO model comparison. Bars represent individual subject LOO scores for Bayes, relative to Fixed. Negative (leftward) values indicate that, for that subject, Bayes had a higher (better) LOO score than Fixed. Blue lines and shaded regions represent, respectively, medians and 95% CI of bootstrapped mean LOO differences across subjects. These values are equal to the summed LOO differences reported in the text divided by the number of subjects.

Although Bayes fits better than Fixed, it still shows systematic deviations from the data, especially at high reliabilities. (Because we fit our models to all of the raw data and because boundary parameters are shared across all reliability levels, the fit to high-reliability trials is constrained by the fit to low-reliability trials.)

#### Noisy log posterior ratio

To see if we could improve Bayes’s fit, we tried a version that included decision noise, i.e. noise on the log posterior ratio *d*. We assumed that this noise takes the form of additive zero-mean Gaussian noise with s.d. *σ*_*d*_. This is almost equivalent to the probability of a response being a logistic (softmax) function of *d*^36^. Adding *d* noise improves the Bayesian model fit by 804 [510, 1134] (Table S1).

For the rest of the reported fits to behavior, we will only consider this version of Bayes with *d* noise, and will refer to this model as Bayes-*d*N. We will refer to Bayes-*d*N, Fixed, Orientation Estimation, Linear Neural, Lin, and Quad, when fitted jointly to category and confidence data from Tasks A and B, as our core models.

#### Heuristic models

Orientation Estimation performs worse than Bayes-*d*N by 2041 [385, 3623] (Figure 5, second row). The intuition for one way that this model fails is as follows: at low levels of reliability, the MAP estimate is heavily influenced by the prior and tends to be very close to the prior mean (0°). This explains why, in Task B, there is a bias towards reporting “high confidence, category 1” at low reliability. Linear Neural performs about as well as Bayes-*d*N, with summed LOO differences of 1188 [-588, 2704], and the fits to the summary statistics are qualitatively poor (Figure 5, third row).

**Figure 5:**
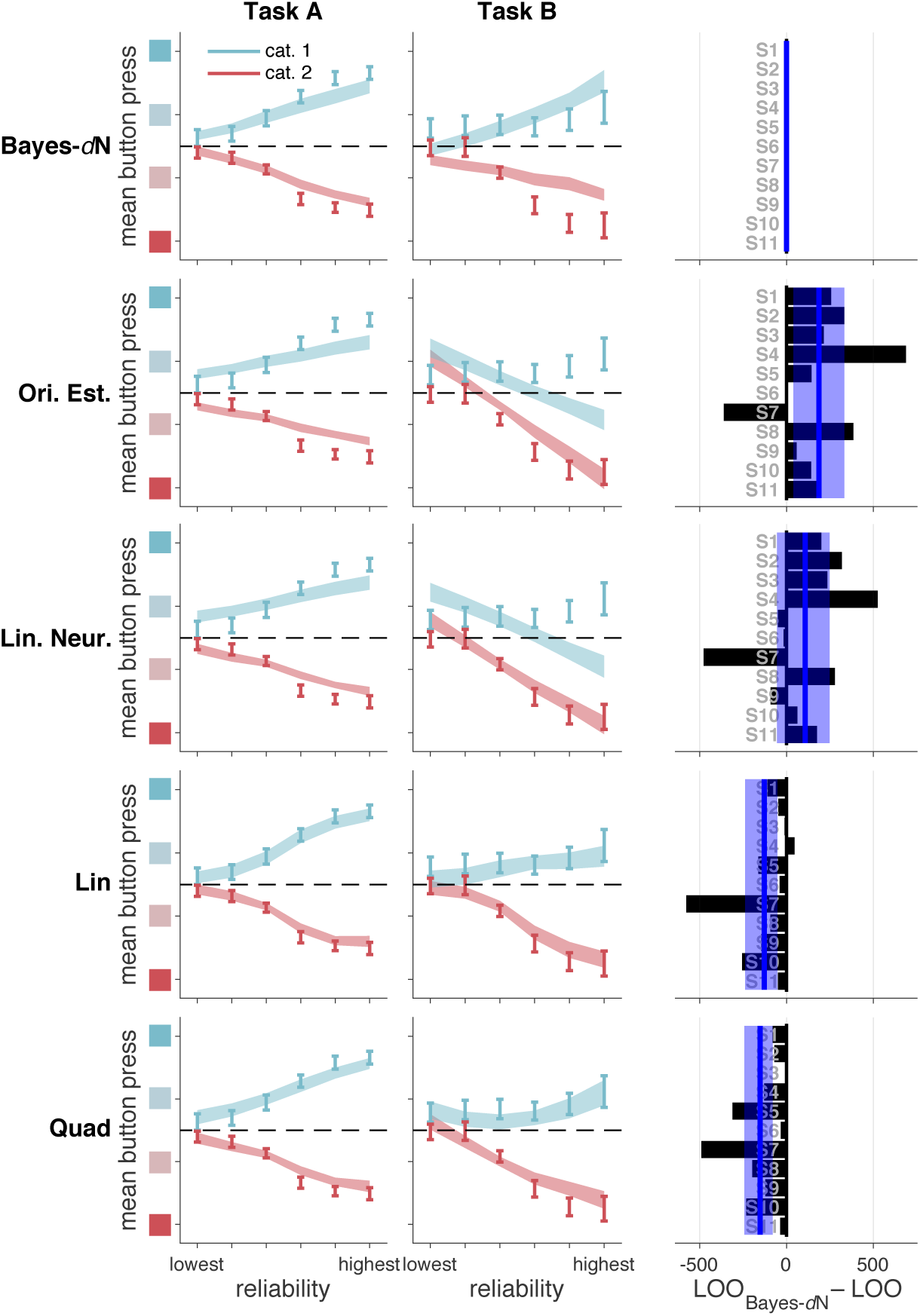
Model fits and model comparison for Bayes-*d*N and heuristic models. In both tasks, Bayes-*d*N fails to describe the data at high reliabilities; Lin and Quad provides a good fit at most reliabilities. Left and middle columns: as in Figure 4. Right column: bars represent individual subject LOO scores for each model, relative to Bayes-*d*N. Negative (leftward) values indicate that, for that subject, the model in the corresponding row had a higher (better) LOO score than Bayes-*d*N. Blue lines and shaded regions: as in Figure 4.

Finally, Lin and Quad outperform Bayes-*d*N by 1398 [571, 2644] and 1667 [858, 2698], respectively. Both models provide qualitatively better fits, especially at high reliabilities (compare Figure 5, first row, to Figure 5, fourth and fifth rows), and strongly tilted orientations (compare Figure S15n,o to Figure S19n,o and Figure 3n,o).

We summarize the performance of our core models in Figure 6. Noting that a LOO difference of more than 5 is considered to be very strong evidence^33^, the heuristic models Lin and Quad perform much better than Bayes-*d*N. Furthermore, we can decisively rule out Fixed. We will now describe variants of our core models.

**Figure 6:**
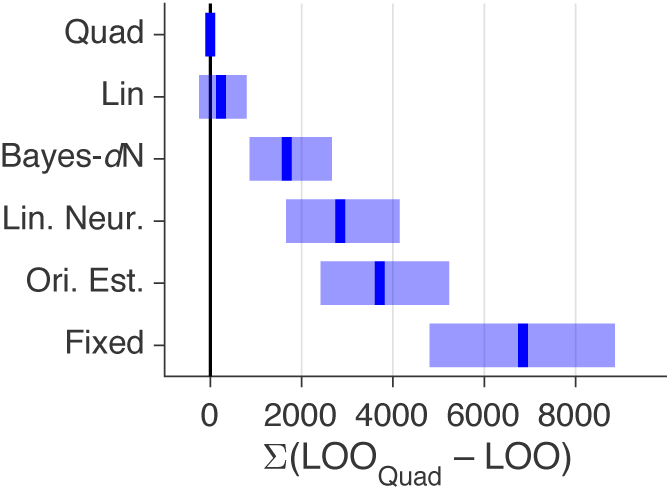
Comparison of core models, experiment 1. Models were fit jointly to Task A and B category and confidence responses. Blue lines and shaded regions represent, respectively, medians and 95% CI of bootstrapped summed LOO differences across subjects. LOO differences for these and other models are shown in Figure S6a.

#### Non-parametric relationship between reliability and *σ*

One potential criticism of our fitting procedure is that we assumed a parameterized relationship between reliability and *σ* (Supplementary Information). To see if our results were dependent on that assumption, we modified the models such that *σ* was non-parametric (i.e., there was a free parameter for *σ* at each level of reliability). With this feature added to our core models, Quad still fits better than Bayes-*d*N by 1676 [839, 2730] and it fits better than Fixed by 6097 [4323, 7901] (Table S1). This feature improved Quad’s performance by 325 [141, 535]. For the rest of this paper, we will only report the fits of Bayes-*d*N, the best-fitting non-Bayesian model, and Fixed. See supplementary figures and tables for all other model fits.

#### Incorrect assumptions about the generative model

Suboptimal behavior can be produced by optimal inference using incorrect generative models, a phenomenon known as “model mismatch”^2,10,56^. Up to now, Bayes-*d*N has assumed that observers have accurate knowledge of the parameters of the generative model. To test whether this assumption prevents Bayes-*d*N from fitting the data well, we tested a series of Bayesian models in which the observer has inaccurate knowledge of the generative model.

Bayes-*d*N assumed that, because subjects were well trained, they knew the true values of *σ*_*C*_, *σ*_1_, and *σ*_2_, the standard deviations of the stimulus distributions. We tested a model in which these values were free parameters, rather than fixed to the true value. We would expect these free parameters to improve the fit of Bayes-*d*N in the case where subjects were not trained enough to sufficiently learn the stimulus distributions. This feature improves Bayes-*d*N’s fit by 908 [318, 1661], but it still underperforms Quad by 768 [399, 1144] (Table S1).

Previous models also assumed that subjects had full knowledge of their own measurement noise; the *σ* used in the computation of *d* was identical to the *σ* that determined their measurement noise. We tested models in which we fit *σ*_measurement_ and *σ*_inference_ as two independent functions of reliability^2^. This feature improves Bayes-*d*N’s fit by 1310 [580, 2175], but it still underperforms Quad by 362 [162, 602] (Table S1).

#### Weighted average of precision and perceived probability of being correct

A recent paper^54^ proposed that confidence is a weighted average of a function of variance, such as _*σ*_^1^_2_, and the perceived probability of being correct (incidentally, under a non-Bayesian decision rule). We tested such a model (using a Bayesian decision rule), which fits better than Fixed by 3059 [758, 5528] but still underperforms Lin by 3478 [2211, 5020] (Table S1).

#### Separate fits to Tasks A and B

In order to determine whether model rankings were primarily due to differences in one of the two tasks, we fit our models to each task individually. In Task A, Quad fits better than Bayes-*d*N by 581 [278, 938], and better than Fixed by 3534 [2529, 4552] (Figure S7 and Table S2). In Task B, Quad fits better than Bayes-*d*N by 978 [406, 1756] and fits better than Fixed by 3234 [2099, 4390] (Figure S8 and Table S3).

#### Fits to category choice data only

In order to see whether our results were peculiar to combined category and confidence responses, we fit our models to the category choices only. Lin fits better than Bayes-*d*N by 595 [311, 927] and fits better than Fixed by 1690 [976, 2534] (Figure S9 and Table S4).

#### Fits to Task B only, with noise parameters fitted from Task A

To confirm that the fitted values of sensory uncertainty in the probabilistic models are meaningful, we treated Task A as an independent experiment to measure subjects’ sensory noise. The category choice data from Task A can be used to determine the four uncertainty parameters. We fit Fixed with a decision boundary of 0° (equivalent to a Bayesian choice model with no prior), using maximum likelihood estimation. We fixed these parameters and used them to fit our models to Task B category and confidence responses. Lin fits better than Bayes-*d*N by 1773 [451, 2845] and fits better than Fixed by 5016 [3090, 6727] (Figure S10 and Table S5).

#### Separate category and confidence responses (experiment 2)

There has been some recent debate as to whether it is more appropriate to collect choice and confidence with a single motor response (as described above) or with separate responses^38,53,69,73^. Aitchison et al.^5^ found that confidence appears more Bayesian when subjects use separate responses. To confirm this, we ran a second experiment in which subjects chose a category by pressing one of two buttons, then reported confidence by pressing one of four buttons. Aitchison et al.^5^ also provided correctness feedback on every trial; in order to ensure that we could compare our results to theirs, we also provided correctness feedback in this experiment, even though this manipulation was not of primary interest. After fitting our core models, our results did not differ substantially from experiment 1: Lin fits better than Bayes-*d*N by 396 [186, 622] and fits better than Fixed by 2095 [1344, 2889] (Figure S11 and Table S6).

#### Task B only (experiment 3)

It is possible that subjects behave suboptimally when they have to do multiple tasks in a session; in other words, perhaps one task “corrupts” the other. To explore this possibility, we ran an an experiment in which subjects completed Task B only. Quad fits better than Bayes-*d*N by 1361 [777, 2022] and fits better than Fixed by 7326 [4905, 9955] (Figure S12 and Table S7). In experiments 2 and 3, subjects only saw drifting Gabors; we did not use ellipses.

We also fit only the choice data, and found that Lin fits about as well as Bayes-*d*N, with summed LOO differences of 117 [-76, 436] and fits better than Fixed by 1084 [619, 1675] (Figure S13 and Table S8). This approximately replicates our previously published results^63^.

#### Model comparison metric

None of our model comparison results depend on our choice of metric: in all three experiments, model rankings changed negligibly if we used AIC, BIC, AICc, or WAIC instead of LOO (Supplementary Information).

## Discussion

Although people can report subjective feelings of confidence, the computations that produce this feeling are not well understood. It has been proposed that confidence is the observer’s computed posterior probability that a decision is correct^20,34,51,62^. However, this hypothesis has not been fully tested. We carried out a strong test of human confidence reports, using overlapping categories^58^, withholding feedback on testing trials, and varying experimental components such as task, stimulus type, and stimulus reliability^49^. We used model comparison to investigate the computational underpinnings of confidence, fitting a total of 75 models from 6 distinct model families.

Our first finding is that, like the optimal observer, subjects use knowledge of their sensory uncertainty when reporting confidence in a categorical decision; models in which the observer ignores their sensory uncertainty provide a poor fit to the data (Figure 4). Our second finding is that subjects do not appear to use knowledge of their sensory uncertainty in a way that is fully consistent with the Bayesian confidence hypothesis. Instead, heuristic models that approximate Bayesian computation—but do not compute a posterior probability over category—outperform the Bayesian models in two tasks (Figure 5, compare top row to bottom two rows). This result continued to hold after we relaxed assumptions about the relationship between reliability and noise, and about the subject’s knowledge of the generating model. We accounted for the fact that our models had different amounts of flexibility by using a wide array of model comparison metrics and by showing that our models are meaningfully distinguishable (Supplementary Information).

Our conclusions differ from those of some recent experimental findings. Rahnev et al.^64^ reported that subjective decision criteria are fixed across conditions of uncertainty. However, their study did not test models in which the criteria were a function of uncertainty, so they cannot make this conclusion very strongly. Additionally, they elicited visibility ratings rather than confidence ratings; the two prompts are known to produce different behavior^65^. Finally, their results may be specific to the case where sensory uncertainty is a function of attention rather than stimulus reliability.

Like the present study, Aitchison et al.^5^ found evidence that confidence reports may emerge from heuristic computations. However, they sampled stimuli from only a small region of their two-dimensional space, where model predictions may not vary greatly. Therefore, their stimulus set did not allow for the models to be strongly distinguished. Furthermore, although they tested for *Bayesian* computation, they did not test for *probabilistic* computation (whether observers take sensory uncertainty into account on a trial-to-trial basis^46^) as we do here. Such a test requires that the experimenter vary the reliability, not only the value, of the stimulus feature of interest.

Sanders et al.^69^ reported that confidence has a “statistical” nature. However, their experiment was unable to determine whether confidence is Bayesian or not^47^, because the stimuli varied along only one dimension. Aitchison et al.^5^ note that, to distinguish models of confidence, the experimenter must use stimuli that are characterized by two dimensions (e.g., contrast and orientation). This is because, when fitting models that map from an internal variable to an integer confidence rating, it is impossible to distinguish between two internal variables that are monotonically related (in the case of Sanders et al. ^69^, the measurement and the posterior probability of being correct). Therefore, the only alternative model proposed by Sanders et al.^69^ is based on reaction time, rather than on the presented stimuli.

Navajas et al.^54^ suggested that confidence reports are best described as a weighted average of precision and the probability of being correct. However, their model uses the estimated probability of being correct under a non-Bayesian decision rule^4^. They did not show the fit of a Bayesian model, and therefore their study does not constitute a true test of whether confidence is Bayesian. Here, we tested and rejected the hypothesis that confidence is a weighted average of precision and the posterior probability of being correct under a Bayesian decision rule.

In this study, we only considered explicit confidence ratings, which differ from the implicit confidence that can be gathered from nonhuman animals^34^ (e.g., by measuring how frequently they decline to make a difficult choice^37^, or how long they will wait for a reward^35^). It is possible that implicit confidence might be more Bayesian^16^. At a minimum, testing this possibility would require an experiment using implicit confidence that could distinguish the models presented here, which has not been done.

What do our findings tell us about the neural basis of confidence? Previous studies have found that neural activity in some brain areas (e.g., human medial temporal lobe^67^ and prefrontal cortex^24^, monkey lateral intraparietal cortex^37^ and pulvinar^40^, rodent orbitofrontal cortex^35^) is associated with behavioral indicators of confidence, and/or with the distance of a stimulus to a decision boundary. However, such studies mostly used stimuli that vary along a single dimension (e.g., net retinal dot motion energy, mixture of two odors). Because measurement is indistinguishable from the probability of being correct in these classes of tasks, neural activity associated with confidence may represent either the measurement or the probability of being correct^5^. In addition to the recommendation of Aitchison et al.^5^ to distinguish between these possibilities by varying stimuli along two dimensions, we recommend fitting both Bayesian and non-Bayesian probabilistic models to behavior. In view of the relatively poor performance of the Bayesian models in the present study, the proposal^62^ to correlate behavior and neural activity with predictions of the Bayesian confidence model should be viewed with skepticism.

Our results raise general issues about the status of Bayesian models as descriptions of behavior. First, because it is impossible to exhaustively test all models that might be considered “Bayesian,” we cannot rule out the entire class of models. However, we have tried to alleviate this issue as much as possible by testing a large number of Bayesian models—far more than the number of Bayesian and non-Bayesian models tested in other studies of confidence. Second, Bayesian models are often held in favor for their generalizability; one can determine the performance-maximizing strategy for any task. Although generalizability indeed makes Bayesian models attractive and powerful, we do not believe that this property should override a bad fit.

One could take two different views of our heuristic model results. The first view is that the heuristics should be taken seriously as principled models^27^; here, the challenge is to demonstrate that they describe behavior in a variety of tasks and can be motivated based on underlying principles. The second view is that these are descriptive models simply meant to demonstrate that a simple model can provide a good fit to the data; here, the heuristics are benchmarks for more principled models, and the challenge is to find a principled model that fits the data as well as the heuristics. We lean towards the second view and interpret our results as demonstrating that the Bayesian confidence hypothesis does not describe human confidence reports particularly well.

However, one might still conclude, after examining the fits of the Bayesian model, that the behavior is “approximately Bayesian” rather than “non-Bayesian.” As written, this is a semantic distinction because it relies on one’s definition of “approximate.” However, it can be turned into a more meaningful question: Are the differences between human behavior and Bayesian models accounted for by an unknown principle, such as an ecologically relevant objective function that includes both task performance and biological constraints?

Although there are benefits associated with veridical explicit representations of confidence^8,9,25^, there are also neural constraints that may give rise to non-Bayesian behavior^11,32^. Such constraints include the kinds of operations that neurons can perform, the high energy cost of spiking^6,43^, and the cost of neural wiring length^17,18^. A search for ecologically rational constraints on Bayesian computation benefits from a positive characterization of the deviations from Bayesian computation, in the form of heuristic models such as Lin and Quad. Specifically, one could define neural networks with various combinations of constraints, and train them as if they were psychophysical subjects in our tasks. After training, one could fit behavioral models to them; this approach has already shown that the output from such neural networks is sometimes best described by heuristic models^57^. Using model ranking as a measure of similarity, one could determine which network architecture and training procedure produces confidence behavior that is most similar to that of humans. This could reveal which constraints are responsible for the specific deviations from Bayesian computation that we have observed.

## ACKNOWLEDGEMENTS

The authors would like to thank Luigi Acerbi for helpful ideas and tools related to model fitting and model comparison. We would also like to thank Luigi Acerbi, Rachel N. Denison, Andra Mihali, A. Emin Orhan, Bas van Opheusden, and Aspen H. Yoo for helpful conversations and comments about the manuscript. This material was based upon work supported by the National Science Foundation Graduate Research Fellowship under Grant No. DGE-1342536.

## AUTHOR CONTRIBUTIONS

W.T.A. and W.J.M. designed the experiments, analyzed the data, and wrote the manuscript. W.T.A. performed the experiments.

## DATA AND CODE AVAILABILITY

All data and code used for running experiments, model fitting, and plotting is available at GitHub/wtadler/confidence.

## Supplementary Information

### 1 Experiment 1

#### 1.1 Subjects

11 subjects (2 male), aged 20–42, participated in the experiment. Subjects received $10 per 40-60 minute session, plus a completion bonus of $15. The experiments were approved by the University Committee on Activities Involving Human Subjects of New York University. Informed consent was given by each subject before the experiment. All subjects were naÏve to the purpose of the experiment. No subjects were fellow scientists.

#### 1.2 Apparatus and stimuli

*Apparatus.* Subjects were seated in a dark room, at a viewing distance of 32 cm from the screen, with their chin in a chinrest. Stimuli were presented on a gamma-corrected 60 Hz 9.7-inch 2048-by-1536 display. The display (LG LP097QX1-SPA2) was the same as that used in the 2013 iPad Air (Apple); we chose it for its high pixel density (264 pixels/inch). The display was connected to a Windows desktop PC using the Psychophysics Toolbox extensions^12,60^ for MATLAB (Mathworks).

*Stimuli.* The background was mid-level gray (199 cd/m^2^). The stimulus was either a drifting Gabor (Subjects 3, 6, 8, 9, 10, and 11) or an ellipse (Subjects 1, 2, 4, 5, and 7). The Gabor had a peak luminance of 398 cd/m^2^ at 100% contrast, a spatial frequency of 0.5 cycles per degrees of visual angle (dva), a speed of 6 cycles per second, a Gaussian envelope with a standard deviation of 1.2 dva, and a randomized starting phase. Each ellipse had a total area of 2.4 dva^2^, and was black (0.01 cd/m^2^). We varied the contrast of the Gabor and the elongation (eccentricity) of the ellipse (Section 1.3).

*Categories.* In Task A, stimulus orientations were drawn from Gaussian distributions with means *μ*_1_ = −4° (category 1) and *μ*_2_ = 4° (category 2) and standard deviations *σ*_1_ = *σ*_2_ = 5°. In Task B, stimulus orientations were drawn from Gaussian distributions with means *μ*_1_ = *μ*_2_ = 0°, and standard deviations *σ*_1_ = 3° (category 1) and *σ*_2_ = 12° (category 2) (Figure 1b). We chose these category means and standard deviations such that the accuracy of an optimal observer would be around 80%.

#### 1.3 Procedure

Each subject completed 5 sessions. Each session consisted of two parts; the subject did Task A in the first part, followed by Task B in the second part, or vice versa (chosen randomly each session). Each part started with instruction and was followed by alternating blocks of 96 category training trials and 144 testing trials, for a total of three blocks of each type, with a block of 24 confidence training trials immediately after the first category training block. Combining all sessions and both tasks, each subject completed 2880 category training trials, 240 confidence training trials, and 4320 testing trials; we did not analyze category training or confidence training trials.

*Instruction.* At the start of each part of a session, subjects were shown 30 (72 in the first session) exemplar stimuli from each category. Additionally, we provided them with a printed graphic similar to Figure 1b, and explained how the stimuli were generated from distributions. We answered any questions.

*Category training.* To ensure that subjects knew the stimulus distributions well, we gave them extensive category training. Each trial proceeded as follows (Figure 1a): Subjects fixated on a central cross for 1 s.

Category 1 or category 2 was selected with equal probability. The stimulus orientation was drawn from the corresponding stimulus distribution (Figure 1b). Gabors had 100% contrast, and ellipses had 0.95 eccentricity (elongation). The stimulus appeared at fixation for 300 ms, replacing the fixation cross. Subjects were asked to report category 1 or category 2 by pressing a button with their left or right index finger, respectively. Subjects were able to respond immediately after the offset of the stimulus, at which point verbal correctness feedback was displayed for 1.1 s. The fixation cross then reappeared.

*Confidence training.* To familiarize subjects with the button mappings, they completed a short confidence training black at the start of every task. We told subjects that in this block, it would be harder to tell what the stimulus orientation was, there would be no correctness feedback, and they would be reporting their confidence on each trial in addition to their category choice. We provided them with a printed graphic similar to the buttons pictured in Figure 1a, indicating that they had to press one of eight buttons to indicate both category choice and confidence level, the latter on a 4-point scale. The confidence levels were labeled as “very high,” “somewhat high,” “somewhat low,” and “very low.” Gabors had 0.4%, 0.8%, 1.7%, 3.3%, 6.7%, or 13.5% contrast, and ellipses had 0.15, 0.28, 0.41, 0.54, 0.67, or 0.8 eccentricity, chosen randomly with equal probability on each trial (Figure 1c). Stimuli were only displayed for 50 ms. Trial-to-trial feedback consisted only of a message telling them which category and confidence level they had reported. Other than these changes, the trial procedure was the same as in category training.

Subjects were not instructed to use the full range of confidence reports^69^, as that might have biased them away from reporting what felt most natural. Instead, they were simply asked to be “as accurate as possible in reporting their confidence” on each trial.

*Testing*. The trial procedure in testing blocks was the same as in confidence training blocks, except that trial-to-trial feedback was completely withheld. At the end of each block, subjects were required to take at least a 30 s break. During the break, they were shown the percentage of trials that they had correctly categorized. Subjects were also shown a list of the top 10 block scores (across all subjects, indicated by initials) for the task they had just done. This was intended to motivate subjects to score highly, and to reassure them that their scores were normal, since it is rare to score above 80% on a block.

#### 1.4 Descriptive statistics

Since our models do not include any learning effects, we wanted to ensure that task performance was stable. For all tasks and experiments, we found no evidence that performance changed significantly as a function of the number of trials. For each experiment and task (the 5 lines in Figure S1), we fit a logistic regression to the binary correctness data for each subject, obtaining a set of slope coefficients. We then used a t-test to determine whether these sets of coefficients differed significantly from zero. In no group did the slopes differ significantly from zero; across all 5 groupings the minimum *p*-value was 0.077 (Task A, experiment 2), which would not be significant even before correcting for multiple comparisons.

**Figure S1:**
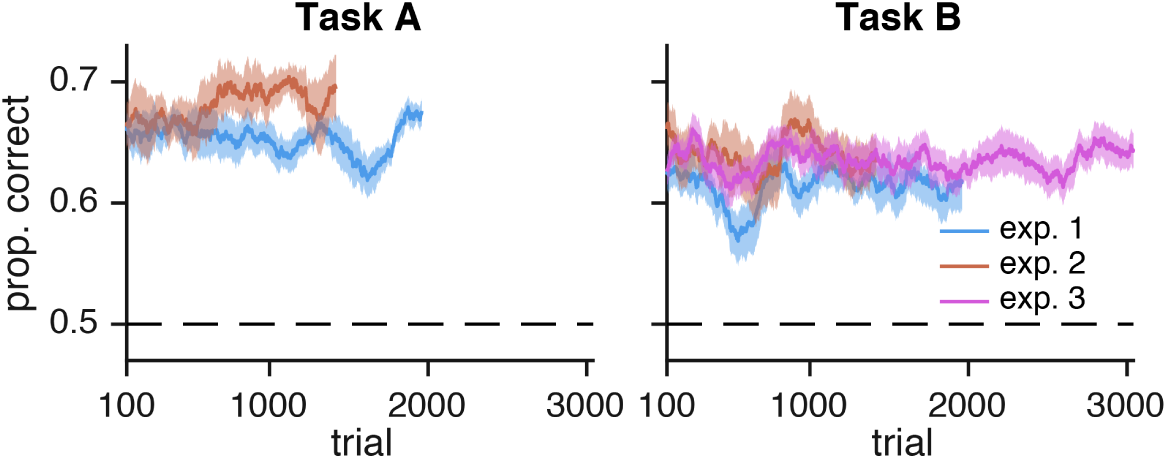
Performance as a function of number of trials, for both tasks and for all experiments. Performance was computed as a moving average over test trials (200 trials wide). Shaded regions represent ±1 s.e.m. over subjects. Performance did not change significantly over the course of each experiment.

##### 1.4.1 Experiment 1

The following statistical differences were assessed using repeated-measures ANOVA.

In Task A, there was a significant effect of true category on category choice (*F*_1,10_ = 285*p* < 10^−7^). There was no main effect of reliability, which took 6 levels of contrast or ellipse elongation, on category choice (*F*_5,50_ = 0.27, p = 0.88). In other words, subjects were not significantly biased to respond with a particular category at low reliabilities. There was a significant interaction between reliability and true category, which is to be expected (*F*_5,50_ = 59.6,*p* < 10^−15^) (Figure 3a).

In Task B, there was again a significant effect of true category on category choice (*F*_1,10_ = 78.3, *p* < 10^−5^). There was no main effect of reliability (*F*_5,50_ = 2.93, *p* = 0.051). There was again a significant interaction between reliability and true category (*F*_5,50_ = 28, *p* < 10^−12^) (Figure 3b).

In Task A, there was a significant effect of true category on response (*F*_1,10_ = 136,*p* < 10^−6^). There was no main effect of reliability (*F*_5,50_ = 0.61, *p* = 0.642). There was a significant interaction between reliability and true category (*F*_5,50_ = 58.7, *p* < 10^−13^) (Figure 3c).

In Task B, there was a significant effect of true category on response (*F*_1,10_ = 54.2,*p* < 10^−6^). There was a significant effect of reliability (*F*_5,50_ = 4.84, *p* = 0.0128). There was a significant interaction between reliability and true category (*F*_5,50_ = 29.2, *p* < 10^−8^) (Figure 3d).

In Task A, there was a main effect of confidence on the proportion of reports (*F*_3,30_ = 7.75,*p* < 10^−3^); low-confidence reports were more frequent than high-confidence reports. There was no significant effect of true category (*F*_1,10_ = 0.784,*p* = 0.397) and no interaction between confidence and category on proportion of responses (*F*_3,30_ = 1.45, *p* = 0.25) (Figure 3e).

In Task B, there was a main effect of confidence on the proportion of reports (*F*_3,30_ = 4.36,*p* = 0.012). There was no significant effect of category (*F*_1,10_ = 0.22,*p* = 0.64), although there was an interaction between confidence and category (*F*_3,30_ = 8.37, *p* = 0.003). This is likely because for task B, category 2 has a higher proportion of “easy” stimuli (Figure 3f).

In both tasks, reported confidence had a significant effect on performance (*F*_3,30_ = 36.9,*p* < 10^−3^). Task also had a significant effect on performance (*F*_1,10_ = 20.1,*p* = 0.001); although we chose the category parameters such that the performance of the optimal observer is matched, subjects were significantly better at Task A. There was no interaction between task and confidence (*F*_3,30_ = 0.878,*p* = 0.436) (Figure 3g).

Figure 3l,m shows psychometric choice curves for both tasks, at all 6 levels of reliability. Each point represents roughly the same number of trials.

Figure 3n,o shows a similar set of psychometric curves. These curves differ from Figure 3l,m in that they represent the mean button press rather than mean category choice.

In Task A (Figure 3l,n), mean category choice and mean button press depend monotonically on orientation, with a slope that increases with reliability. In Task B (Figure 3m,o), the mean category choice and mean button press tends towards category 1 when stimulus orientation is near horizontal, and tends towards category 2 when orientation is strongly tilted; this reflects the stimulus distributions.

*Effect of stimulus type on results: Gabor vs. ellipse*. Since some subjects only saw Gabors and some only saw ellipses, we used Spearman’s rank correlation coefficient to measure the similarity of the two groups’ model rankings. Spearman’s rank correlation coefficient between Gabor and ellipse subjects for the summed LOO scores of the model groupings in Figure 6 and Figure S6 was 0.952 and 0.944, respectively (a value of 1 would indicate identical rankings). In both model groupings, the identities of the lowest-and highest-ranked models were the same for both Gabor and ellipse subjects. This indicates that the choice of stimulus type did not have a systematic effect on model rankings.

## 2 Experiment 2: Separate category and confidence responses and testing feedback

This control experiment was identical to experiment 1 except for the following modifications:

- Subjects first reported choice by pressing one of two buttons with their left hand, and then reported confidence by pressing one of four buttons with their right hand.
- Subjects reported confidence in category training blocks, and received correctness feedback after reporting confidence.
- There were no confidence training blocks.
- In testing blocks, subjects received correctness feedback after each trial.
- Subjects completed a total of 3240 testing trials.
- 8 subjects (0 male), aged 19-23, participated. None were participants in experiment 1, and again, none were fellow scientists.
- Drifting Gabors were used; no subjects saw ellipses.

## 3 Experiment 3: Task B only

This experiment was identical to experiment 1 except for the following modifications:

- Subjects completed blocks of Task B only.
- Subjects completed a total of 3240 testing trials.
- 15 subjects (7 female), aged 19-30, participated. None were participants in experiments 1 or 2.
- Drifting Gabors were used; no subjects saw ellipses.

## 4 Modeling

### 4.1 Measurement noise

For models (such as our core models) where the relationship between reliability (i.e., contrast or ellipse eccentricity) and noise was parametric, we assumed a power law relationship between reliability *c* and measurement noise variance *σ*^2^: *σ*^2^(*c*) = γ + *αc*^−β^. We have previously^63^ used this power law relationship because it encompasses a large family of monotonically decreasing relationships using only three parameters. The relationship is also consistent with a form of the Naka-Rushton function^19,52^ commonly used to describe the mapping from reliability to neural gain 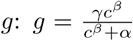. The power law relationship then holds under the assumption that measurement noise variance is inversely proportional to gain^48^.

For all models except the Bayesian model with additive precision, we assumed additive orientation-dependent noise in the form of a rectified 2-cycle sinusoid, accounting for the finding that measurement noise is higher at non-cardinal orientations^28^. The measurement noise s.d. comes out to

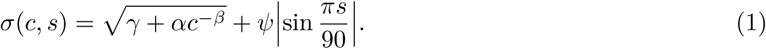

### 4.2 Response probability

We coded all responses as *r* ∈ {1,2, …, 8}, with each value indicating category and confidence. For all models except the Linear Neural model, the probability of a single trial *i* is equal to the probability mass of the measurement distribution 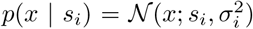 (i.e., a normal distribution over *x* with mean *s*_*i*_ and variance 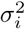) in a range corresponding to the subject’s response *r*_*i*_. Because we only use a small range of orientations, we can safely approximate measurement noise as a normal distribution rather than a Von Mises distribution. We find the boundaries (*b*_*r*_*i*_−1_(*σ*_*i*_), *b*_*r*_*i*__(*σ*_*i*_)) in measurement space, as defined by the fitting model and parameters *θ* and then compute the probability mass of the measurement distribution between the boundaries:

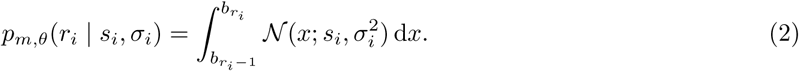

For Task A, *b*_0_ = -∞° and *b*_8_ = ∞°. For Task B, *b*_0_ = 0° and *b*_8_ = ∞°; since the task is symmetric around 0°, we only use |s| in our computation of the log likelihood.

To obtain the log likelihood of the dataset, given a model with parameters *θ* we compute the sum of the log probability for every trial *i*, where *t* is the total number of trials:

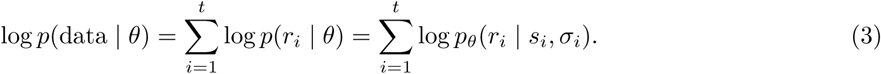

### 4.3 Model specification

#### 4.3.1 Bayesian

*Derivation of d_A_ and _B_*. The log posterior ratio *d* is equivalent to the log likelihood ratio plus the log prior ratio:

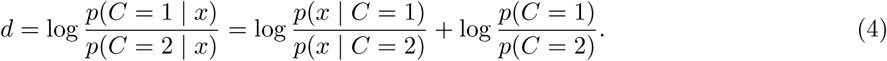

To get *d*_*A*_ and *d*_*B*_, we need to find the task-specific expressions for *p*(*x* | *C*). The observer knows that the measurement *x* is caused by the stimulus *s*, but has no knowledge of *s*. Therefore, the optimal observer marginalizes over *s*:

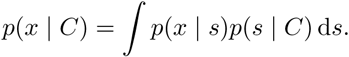

We substitute the expressions for the noise distribution and the stimulus distribution, and evaluate the integral:

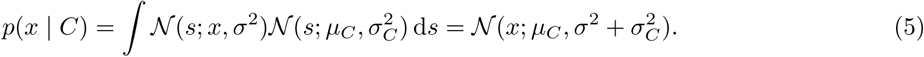

Plugging the task-and category-specific *μ_C_* and *σC* into Equation (5), and substituting the resulting expression back into Equation (4), we get:

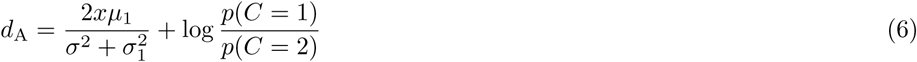

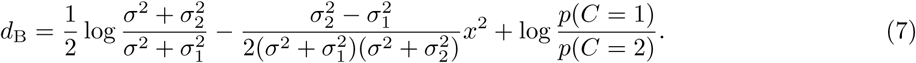

The 8 possible category and confidence responses are determined by comparing the log posterior ratio *d* to a set of decision boundaries **k** = (*k*_0_, *k*_1_, …, *k*_g_). *k*_4_ is equal to the log prior ratio 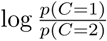, which functions as the boundary on *d* between the 4 category 1 responses and the 4 category 2 responses; *k*_0_ is the only boundary parameter in models of category choice (and not confidence). *k*_0_ is fixed at-∞ and *k*_8_ is fixed at ∞. In all models, the observer chooses category 1 when *d* is positive.

Because the decision boundaries are free parameters, our models effectively include a large family of possible cost functions. A different cost function would be equivalent to a rescaling of the confidence boundaries **k**. To see this, it is probably easiest to consider category choice alone; there, asymmetric costs for getting either category wrong would translate into a different value of *k*j, the category decision boundary (i.e., the observer’s prior over category). For us, this boundary (like all other boundaries) is a free parameter.

The posterior probability of category 1 can be written as as 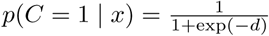.

*Levels of strength.* The Bayesian model is unique in that it is possible to formulate a principled version with relatively few boundary parameters. In principle, it is possible that such a model could perform better than more flexible models, if those models are overfitting. We formulated several levels of strength of the BCH, with weaker versions having fewer assumptions and more sets of mappings between the posterior probability of being correct and the confidence report (Figure S2). In the *ultrastrong BCH*, confidence is a function solely of the posterior probability of the chosen category. In the *strong BCH*, it is additionally a function of the current task.

Most studies cannot distinguish between the ultrastrong and strong BCH because they test subjects in only one task. Furthermore, the weak BCH is only justifiable in tasks where the categories have different distributions of the posterior probability of being correct; the subject may then rescale their mappings between the posterior and their confidence. Here, one can see that Task B has this feature by observing that, in the bottom row of Figure S2, the distributions of posterior probabilities are different for the two categories). Most experimental tasks are like Task A, where the distributions are identical. We compared Bayesian models (Bayes_Ultrastrong_, Bayes_Strong_) corresponding to each of these versions of the BCH.

In Bayes_UltraStrong_, **k** is symmetric across *k*_4_: *k*_4+j_ - k_4_ = k_4_ - k_4-j_ for *j* ∈ {1,2,3}. Furthermore, in Bayes_Ultrastrong_, **k**_A_ = **k**_B_. So Bayes_Ultrastrong_ has a total of 4 free boundary parameters: *k*_1_,*k*_2_,*k*_3_,*k*_4_. Bayes_Ultrastrong_ consists of the observer determining the response by comparing *d*_*A*_ and *d*_*B*_ to a single symmetric set of boundaries (Figure S2, left column).

Bayes_Strong_ is identical to Bayes_Ultrastr_ except that **kA** is allowed to differ from **kB**. So Bayes_Strong_ has a total of 8 free boundary parameters: *k* _1_A, *k*2A, *k*3A, *k*4A, *k* _1_B, *k*2B, *k*3B, *k*4B. Bayes_Strong_ consists of the observer determining the response by comparing *d_A_* to a symmetric set of boundaries, and *d*_B_ to a different symmetric set of boundaries (Figure S2, middle column).

Bayes_Weak_ is identical to Bayes_Strong_ except that symmetry is not enforced for **kB**. So Bayes_Weak_ has a total of 11 free boundary parameters: *k1_A_, k2A, k3_A_, k*4A, *k* _1_B, *k*2B, *k3B, k*4B, *k*5B, *k*6B, *k*7B. Bayes_Weak_ consists of the observer comparing *d_A_* to a symmetric set of boundaries, and *d*_B_ to a different non-symmetric set of boundaries (Figure S2, right column).

We did not include Bayes_Strong_ and Bayes_Ultrastrong_ in the core models reported in the main text, because Bayes_Weak_ provided a much better fit to the data. Because it was not necessary in the main text to distinguish the three strengths of Bayesian models, we refer to Bayes_Weak_ there simply as Bayes. However, we do include Bayes_Strong_ and Bayes_Ultrastrong_ in our model recovery analysis (Section 4.9) and in our supplemental model comparison tables.

*Decision boundaries.* In the Bayesian models without *d* noise, we translate boundary parameters **k** to measurement boundaries **b** corresponding to fitted noise levels *σ.* To do this, we use the parameters **k** as the left-hand side of Equations (6) and (7) and solve for *x* at the fitted levels of *σ*. These values were used as the measurement boundaries **b**(*σ*).

**Figure S2:**
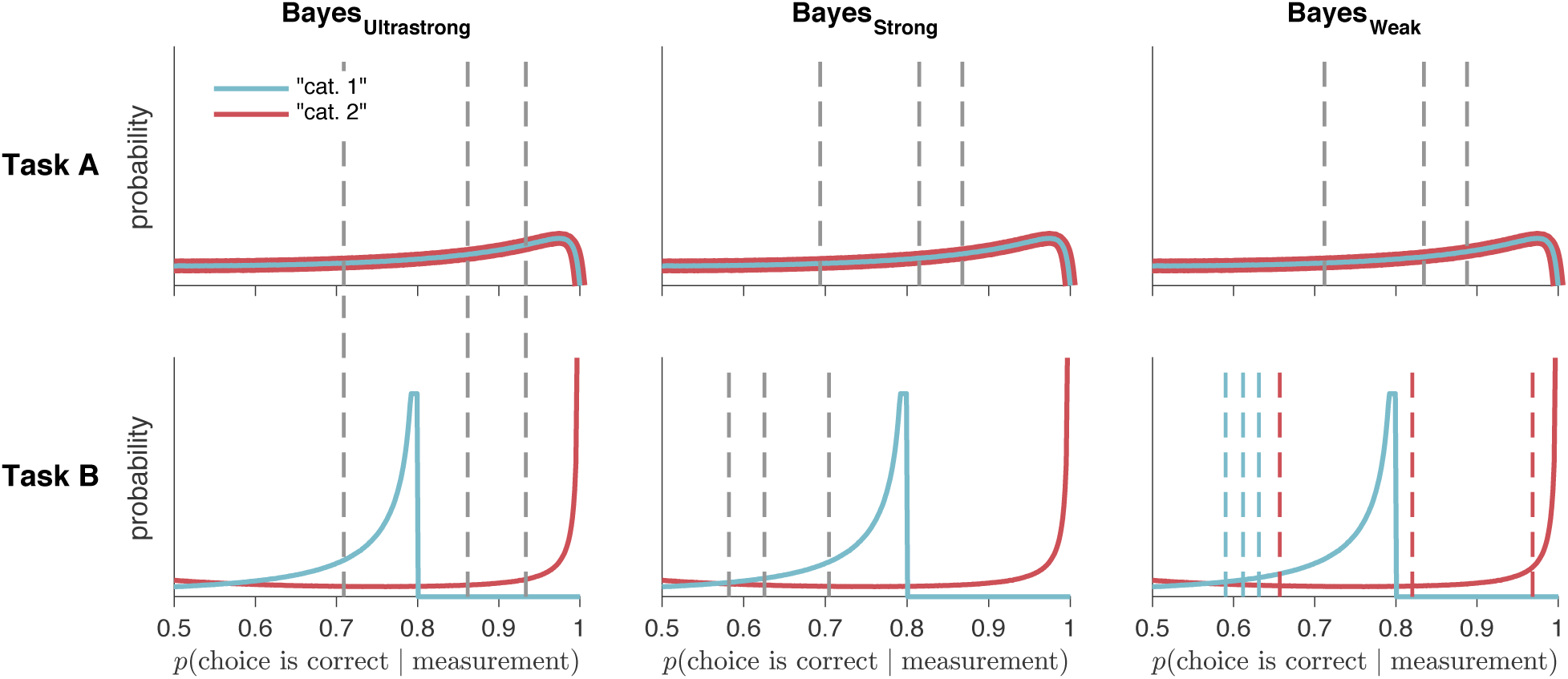
Distributions of posterior probabilities of being correct, with confidence criteria for Bayesian models with three different levels of strength. Solid lines represent the distributions of posterior probabilities for each category and task in the absence of measurement noise. Dashed lines represent confidence criteria, generated from the mean of subject 4’s posterior distribution over parameters. Each model has a different number of sets of mappings between posterior probability and confidence report. In Bayes_Ultrastrong_, there is one set of mappings. In Bayes_Strong_, there is one set for Task A, and another for Task B. In Bayes_Weak_, as in the non-Bayesian models, there is one set for Task A, and one set for each reported category in Task B. Plots were generated from the mean of subject 4’s posterior distribution over parameters as in Figure 2.

In the Bayesian models with *d* noise, we assume that, for each trial, there is an added Gaussian noise term on *d*, *ηd* ~ *p*(*ηd*), where 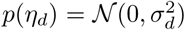, and *σ*_*d*_ is a free parameter. **W**e pre-computed 101 evenly spaced draws of *ηd* and their corresponding probability densities *p*(*ηd*). **W**e used Equations (6) and (7) to compute a lookup table containing the values of *d* as a function of *x, σ*, and *ηd.* **W**e then used linear interpolation to find sets of measurement boundaries **b**(*σ*) corresponding to each draw of *ηd*^1^. We then computed 101 response probabilities for each trial (Section 4.2), one for each draw of *ηd*, and computed the weighted average according to *p*(*ηd*).

#### 4.3.2 Probability correct with additive precision

We tested a model in which the decision variable was a weighted mixture of precision (equivalent in this case to the Fisher information of the measurement variable *x*) and the perceived probability of being correct^54^. In this model, the decision variable is 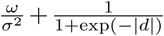, where *ω* is a free parameter. To find the measurement boundaries **b**(*σ*), we substituted Equations (6) and (7) for *d*, and set the whole value equal to parameters **k**, solving for *x* at the fitted levels of *σ*. This model can be considered a hybrid Bayesian-heuristic model. Like Bayes_Ultrastrong_, it has 4 free boundary parameters. Although the model is a hybrid Bayesian-heuristic model, not a strictly Bayesian one, we refer to it as Bayes_Ultrastrong_ + precision in Figure S6 and Table S1.

#### 4.3.3 Fixed

In Fixed, the observer compares the measurement to a set of boundaries that are not dependent on *σ.* We fit free parameters **k**, and use measurement boundaries *b*_*r*_ = *k*_*r*_.

#### 4.3.4 Lin and Quad

In Lin and Quad, the observer compares the measurement to a set of boundaries that are linear or quadratic functions of *σ*. We fit free parameters **k** and **m**, and use measurement boundaries *b_r_(σ) = k_r_ + m_r_σ* (Lin) or *b*_*r*_(σ) = *k*_*r*_ + *m*_*r*_σ^2^ (Quad).

Lin and Quad are each a supermodel of Fixed. In other words, there are parameter settings where Lin and Quad are equivalent to Fixed (although our model comparison methods ensure that the models are still distinguishable, see Section 4.9). Additionally, in Task A, Quad is a supermodel of the Bayesian models without *d* noise.

#### 4.3.5 Orientation Estimation

In Orientation Estimation, the observer uses the mixture of the two stimulus distributions as a prior distribution to compute a maximum a posteriori estimate of the stimulus:

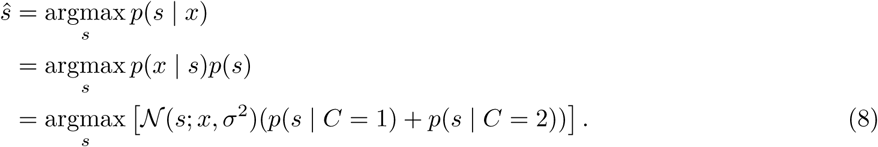

The observer then compares *ŝ* to a set of boundaries **k** to determine category and confidence response.

*Decision boundaries.* To find the decision boundaries in measurement space, we used *gmm1max_n2_fast* from Luigi Acerbi’s gmm1 (github.com/lacerbi/gmm1) 1-D Gaussian mixture model toolbox to solve Equation (8), computing a lookup table containing the value of *s* as a function of *x* and *σ*^2^. We then found, using linear interpolation, the values of *x* corresponding to *σ* and the free parameters **k**. These values were used as the measurement boundaries **b**(*σ*).

#### 4.3.6 Linear Neural

In this section, **r** refers to neural activity, not button responses. This model is different from all other models in that the generative model does not include measurement *x.* The model can be derived as follows.

All neurons have Gaussian tuning curves with variance 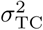 and gain 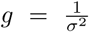. Tuning curve means are contained in the vector of preferred stimuli 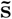. The number of spikes in the population is r ~ Poisson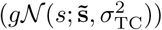. Neural weights are a linear function of the preferred stimuli: 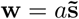.

On each trial, we get some quantity that is a weighted sum of each neuron’s activity, *z* = w · r · 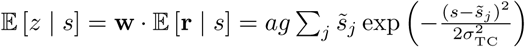.

Rather than sum over all neurons, we assume an infinite number of neurons uniformly spanning all possible preferred stimuli 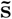. This allows us to replace the sum with an integral. The expected value of *z* is 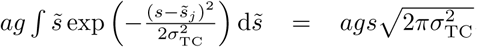. The variance of *z* is 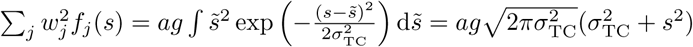.

Now that we have the mean and variance of *z*, we assume that *z* is normally distributed. This is equivalent to assuming that there are a high number of spikes, because the Poisson distribution approximates the normal distribution as the rate parameter becomes high. To compute response probability, we fit neural activity boundaries **k**, and replace Equation (2) with

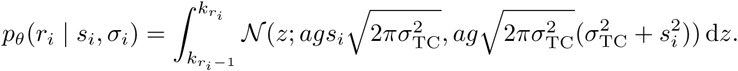

### 4.4 Lapse rates

In confidence and category models, we fit three different types of lapse rate. On each trial, there is some fitted probability of:

- A “full lapse” in which the category report is random, and confidence report is chosen from a distribution over the four levels defined by λ_1_, the probability of a “very low confidence” response, and A4, the probability of a “very high confidence” response, with linear interpolation for the two intermediate levels.
- A “confidence lapse” λ_confidence_ in which the category report is chosen normally, but the confidence report is chosen from a uniform distribution over the four levels.
- A “repeat lapse” λ_repeat_ in which the category and confidence response is simply repeated from the previous trial.

In category choice models, we fit a standard category lapse rate λ, as well as the above “repeat lapse” λ_repeat_.

### 4.5 Parameterization

Because of tradeoffs when directly fitting parameters α, β, γ, we re-parameterized Equation (1) as

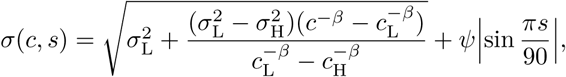

where *C*_L_ and *C*_H_ were the values of the lowest and highest reliabilities used. This way, CTL and CTH were free parameters that determined the s.d. of the measurement distributions for the lowest and highest reliabilities, and β was a free parameter determining the curvature of the function between the two reliabilities. For models where the relationship between reliability and noise was non-parametric, the first term in Equation (1) was replaced with free s.d. parameters (σr_rel. 1_,…, σ_rel. 6_) corresponding to each of the six reliability levels.

For models where subjects had incorrect knowledge about their measurement noise, we fit two sets of uncertainty-related parameters. One set was for the generative noise (used in Equation (2)), and the other set was for the subject’s believed noise (used in Equations (6) to (8)).

All parameters that defined the width of a distribution (e.g., σ_L_, σ_H_, σ_*d*_, σ_rel. 1_, …) were sampled in log-space and exponentiated during the computation of the log likelihood. See *model_parameters.xls* for a complete list of each model’s parameters.

### 4.6 Model fitting

Rather than find a maximum likelihood estimate of the parameters, we sampled from the posterior distribution over parameters, *p*(*θ* | data); this has the advantage of maintaining a measure of uncertainty about the parameters, which can be used both for model comparison and for plotting model fits. We used the log posterior

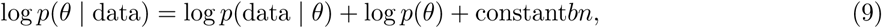

where log*p*(data | *θ*) is given in Equation (3). We assumed a factorized prior over each parameter *j*:

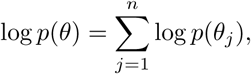

where *j* is the parameter index and *n* is the number of parameters. We took uniform (or, for parameters that were standard deviations, log-uniform) priors over reasonable, sufficiently large ranges^2^, which we chose before fitting any models.

We sampled from the probability distribution using a Markov Chain Monte Carlo (MCMC) method, slice sampling^55^. For each model and dataset combination, we ran between 4 and 7 parallel chains with random starting points. For each chain, we took 40,000 to 600,000 total samples (depending on model computational time) from the posterior distribution over parameters. We discarded the first third of the samples and kept 6,667 of the remaining samples, evenly spaced to reduce autocorrelation. All samples with log posteriors more than 40 below the maximum log posterior were discarded. Marginal probability distributions of the sample log likelihoods were visually checked for convergence across chains. In total we had 842 model and dataset combinations, with a median of 26,668 kept samples (IQR = 13,334).

After sampling, we conducted a visual check to confirm that our parameter ranges were sufficiently large. For each model, we plotted the posterior distribution over parameter values for each subject; an example plot is shown in Figure S3. Visual checks of these plots confirmed that the distributions are unimodal and roughly Gaussian. Visual checks also confirmed that the parameter distributions are well-contained within the chosen parameter ranges, except for the distributions of:

- Lapse rate parameters, which tend to mass around 0, where they are necessarily bounded.
- Log noise parameters, which have a large negative range where they are effectively at zero noise.
- Upper confidence boundary parameters, which become small for subjects who frequently report “high confidence,” or large for subjects who frequently do.

**Figure S3:**
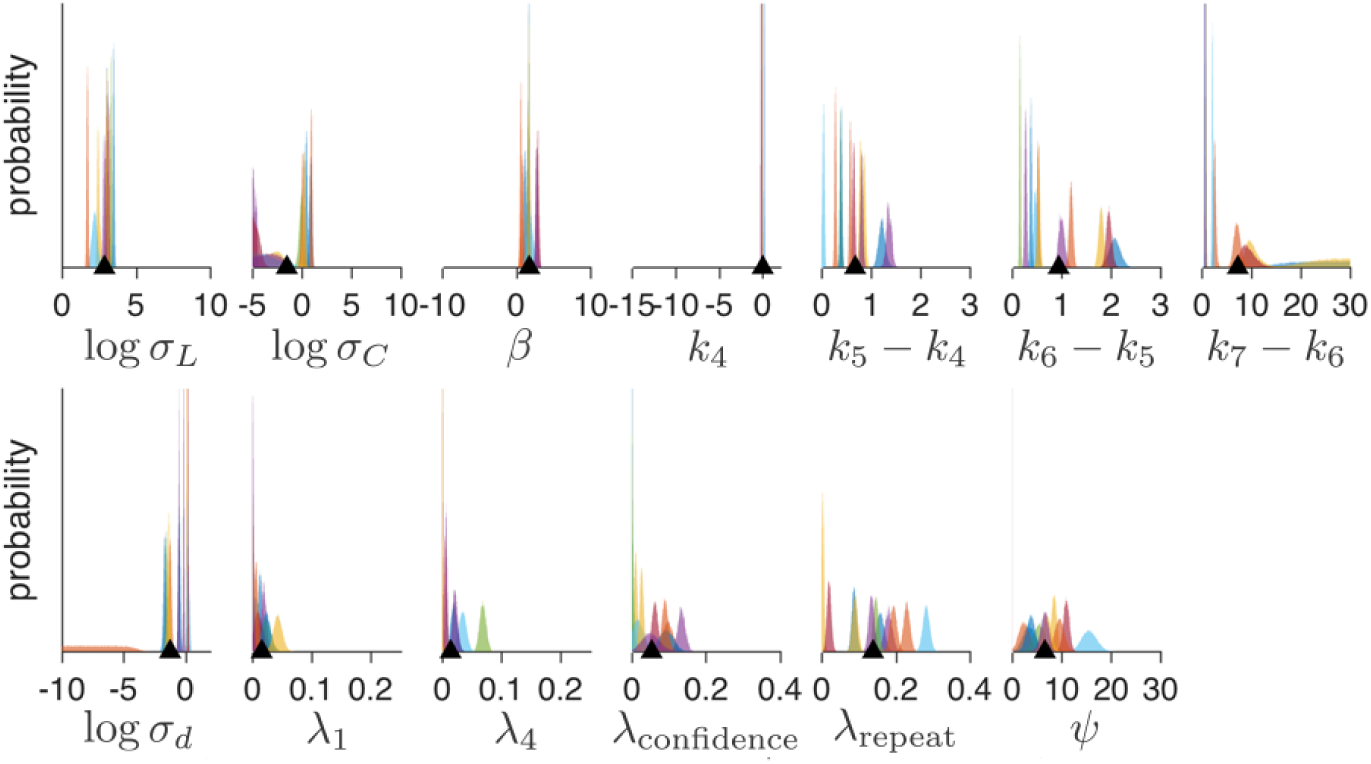
Posterior distributions over parameter values for an example model. Each subplot represents a parameter of the model. Each colored histogram represents the sampled posterior distribution for a parameter and a subject in experiment 1, with colors consistent for each subject. The limits of the x-axis indicates the allowable range for each parameter. Black triangles indicate the overall mean parameter value.

### 4.7 Model comparison

#### 4.7.1 Model groupings

We used 8 groupings of model-subject combinations where it made sense to consider the models as being on equal footing for the purpose of model comparison. The model-subject combinations were grouped by: experiment (which corresponded to subject population), data type (category response only vs. category and confidence response), task type (Task A, B, or both fit jointly). The 8 groupings correspond to Figures S6 to S13 and Tables S1 to S8.

#### 4.7.2 Metric choice

A more complex model is likely to fit a dataset better than a simpler model, even if only by chance. Since we are interested in our models’ predictive accuracy for unobserved data, it is important to choose a metric for model comparison that takes the complexity of the model into account, avoiding the problem of overfitting. Roughly speaking, there are two ways to compare models: information criteria and cross-validation.

Most information criteria (such as AIC, BIC, and AICc) are based on a point estimate for *θ*, typically *θMLE*, the *θ* that maximizes the log likelihood of the dataset (Equation (3)). For instance, AIC adds a correction for the number of parameters *n* to the log likelihood of the dataset: 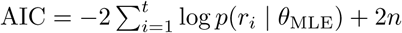.

WAIC is a more Bayesian approach to information criteria that adds a correction for the effective number of parameters^26^. Because WAIC is based on samples from the full posterior of *θ* (Equation (9), typically sampled via MCMC), it takes into account the model’s uncertainty landscape.

Although information criteria are computationally convenient, they are based on asymptotic results and assumptions about the data that may not always hold^26^. An alternative way to estimate predictive accuracy for unobserved data is to cross-validate, fitting the model to training data and evaluating the fit on held out data. Leave-one-out cross-validation is the most thorough way to cross-validate, but is very computationally intensive; it requires that you fit your model *t* times, where *t* is the number of trials. Here we use a method (PSIS-LOO, referred to here simply as LOO) proposed by Vehtari et al.^72^ for approximating leave-one-out cross-validation that, like WAIC, uses samples from the full posterior of *θ*:

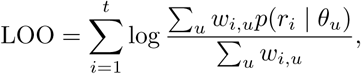

where *θ*_u_ is the *u*-th sampled set of parameters, and *w*_*i,u*_ is the importance weight of trial *i* for sample *u*. Pareto smoothed importance sampling provides an accurate and reliable estimate of the weights. LOO is currently the most accurate approximation of leave-one-out cross-validation^3^. Conveniently, it has a natural diagnostic that allows the user to know when the metric may be inaccurate^72^; we used that diagnostic and confirmed that our use of the metric is justified.

We determined that our results were not dependent on our choice of metric. We computed AIC, BIC, AICc, WAIC, and LOO for all models in the 8 model groupings, multiplying the information criteria by −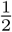 to match the scale of LOO. For AIC, BIC, and AICc, we used the parameter sample with the highest log likelihood as our estimate of *θMLE.* Then we computed Spearman’s rank correlation coefficient for every possible pairwise comparison of model comparison metrics for all model and dataset combinations, producing 80 total values (8 model groupings **x** 10 possible pairwise comparisons of model comparison metrics). All values were greater than 0.998, indicating that, had we used an information criterion instead of LOO, we would not have changed our conclusions. Furthermore, there are no model groupings in which the identities of the lowest-and highest-ranked models are dependent on the choice of metric. The agreement of these metrics strengthens our confidence in our conclusions.

#### 4.7.3 Metric aggregation

*Summed LOO differences.* In all figures where we present model comparison results (e.g., Figure 5, right column), we aggregate LOO scores by the following procedure. Choose a reference model (usually the one with the lowest mean LOO score across subjects). Subtract all LOO scores from the corresponding subject’s score for the reference model; this converts all scores to a LOO “difference from reference” score, with higher scores being worse. Repeat the following standard bootstrap procedure 10,000 times: Choose randomly, with replacement, a group of datasets equal to the total number of unique datasets, and take the sum over subjects of their “difference from reference” scores for each model. Plots indicate the median and 95% CI of these bootstrapped summed “difference from reference” scores. This approach implicitly assumes that all data was generated from the same model.

To confirm that our sample size was large enough to trust our bootstrapped confidence intervals, we bootstrapped our bootstrapping procedure to see how the confidence intervals were affected by the number of subjects *N*. For an example pair of models that we might be interested in comparing, and took the 11 LOO differences between the models, one for each subject in experiment 1. For each *N* between 2 and 11, we took 50 subsamples of our subject LOO differences with replacement; this is akin to running the experiment 50 times for each *N*. For each subsample, we conducted the above bootstrap procedure, which give us a median and 95% CI on the mean of differences. We then plot the mean of these values, with error bars indicating ±1 s.d., at each *N* (Figure S4a). A visual check indicates that the confidence interval appears to converge at about *N* = 9. This indicates that our bootstrapped confidence intervals are trustworthy.

**Figure S4:**
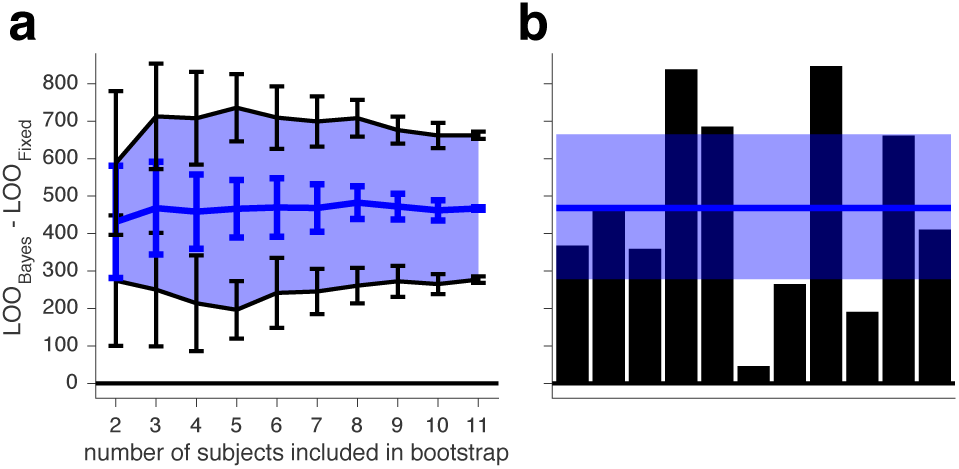
Example analysis of a bootstrapped confidence interval. (**a**) Uncertainty estimates for bootstrapped confidence intervals, as a function of the number of subjects included. Blue line represents the median bootstrapped mean of LOO differences, and black lines indicate the lower and upper bounds of the 95% CI. Error bars represent ±1 s.d. (**b**) For comparison to **a**, the standard style of plot used to show model comparison results (e.g., Figure 4).

*Group level Bayesian model selection.* We also used LOO scores to compute two metrics that allow for model heterogeneity across the group. The first metric was “protected exceedance probability,” the posterior probability that one model occurs more frequently than any other model in the set^66^, above and beyond chance (e.g., Figure S6b). The second was the expected posterior probability that a model generated the data of a randomly chosen dataset^70^ (e.g., Figure S6c). The latter metric assumes a uniform prior over models, which is a function of the total number of datasets. We used the SPM12 (www.fil.ion.ucl.ac.uk/spm) software package to compute these metrics.

In all but one of the 8 model groupings, all three methods of metric aggregation identify the same overall best model. For example, in Figure S6, one model (Quad + non-param. *σ*) has the lowest summed LOO differences, the highest protected exceedance probability, and the highest expected posterior probability.

### 4.8 Visualization of model fits

Model fits were plotted by bootstrapping synthetic group datasets with the following procedure: For each task, model, and subject, we generated 20 synthetic datasets, each using a different set of parameters sampled, without replacement, from the posterior distribution of parameters. Each synthetic dataset was generated using the same stimuli as the ones presented to the real subject. We randomly selected a number of synthetic datasets equal to the number of subjects to create a synthetic group dataset. For each synthetic group dataset, we computed the mean output (e.g., button press, confidence, performance) per bin. We then repeated this 1,000 times and computed the mean and standard deviation of the mean output per bin across all 1,000 synthetic group datasets, which we then plotted as the shaded regions. Therefore, shaded regions represent the mean ±1 s.e.m. of synthetic group datasets.

For plots with orientation on the horizontal axis (e.g., Figure 3j-o), stimulus orientation was binned according to quantiles of the task-dependent stimulus distributions so that each point consisted of roughly the same number of trials. For each task, we took the overall stimulus distribution *p*(*s*) = 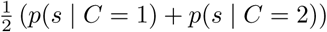 and found bin edges such that the probability mass of *p(s*) was the same in each bin. We then plotted the binned data with linear spacing on the horizontal axis.

### 4.9 Model recovery

We performed a model recovery analysis^71^ to test our ability to distinguish our 6 core models, as well as the 2 stronger versions of the Bayesian model(Section 4.3.1). We generated synthetic datasets from each of the 8 models for both Tasks A and B, using the same sets of stimuli that were originally randomly generated for each of the 11 subjects. To ensure that the statistics of the generated responses were similar to those of the subjects, we generated responses to these stimuli from 4 of the randomly chosen parameter estimates obtained via MCMC sampling (as described in Section 4.6) for each subject and model. In total, we generated 352 datasets (8 generating models × 11 subjects × 4 datasets). We then fit all 8 models to every dataset, using maximum likelihood estimation (MLE) of parameters by an interior-point constrained optimization (MATLAB’s *fmincon*), and computed AIC scores from the resulting fits.

We found that the true generating model was the best-fitting model, on average, in all cases (Figure S5). Overall, AIC “selected” the correct model (i.e., AIC scores were lowest for the model that generated the data) for 86.6% of the datasets, indicating that our models are distinguishable.

**Figure S5:**
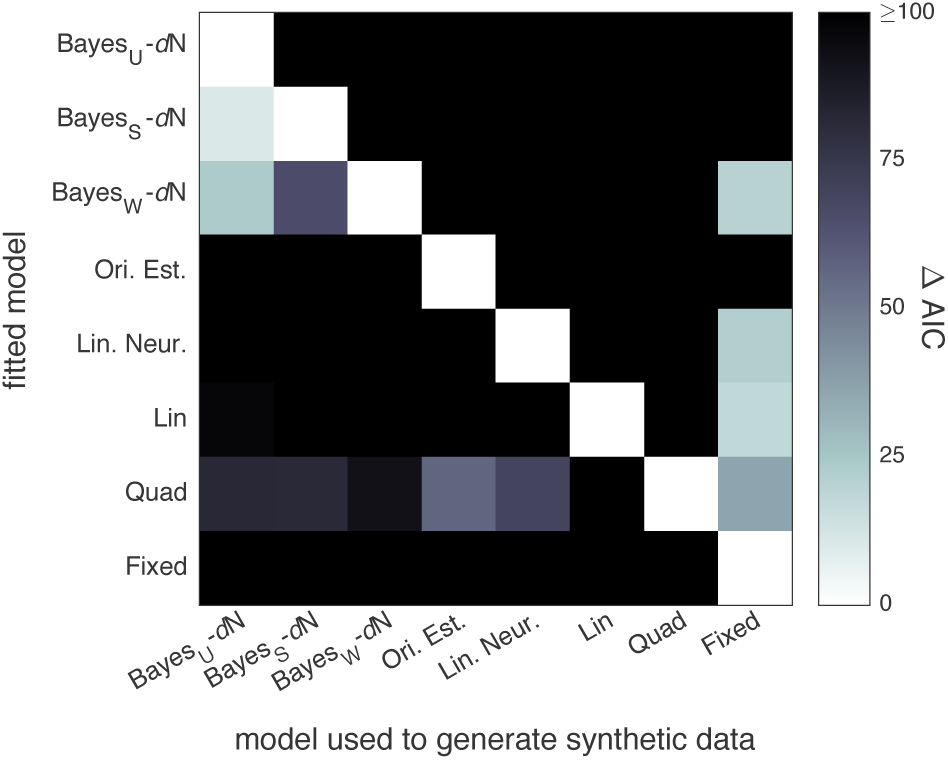
Model recovery analysis. Shade represents the difference between the mean AIC score (across datasets) for each fitted model and for the one with the lowest mean AIC score. White squares indicate the model that had the lowest mean AIC score when fitted to data generated from each model. The squares on the diagonal indicate that the true generating model was the best-fitting model, on average, in all cases.

Ideally, we would have evaluated our model recovery fits using LOO, as we evaluated the fits to human data. However, LOO can only be obtained when fitting with MCMC sampling, which takes orders of magnitudes longer than fitting with MLE. It would be impossible to fit all 352 synthetic datasets in a short amount of time using the same procedure and sampling quality standards described in Section 4.6 (i.e., a large number of samples, across multiple converged chains). Furthermore, we do not believe that our model recovery is dependent on how the models are fit and the fits are evaluated; we found that AIC and LOO scores gave us near-identical model rankings for data from real subjects (Section 4.7.2).

**Figure S6:**
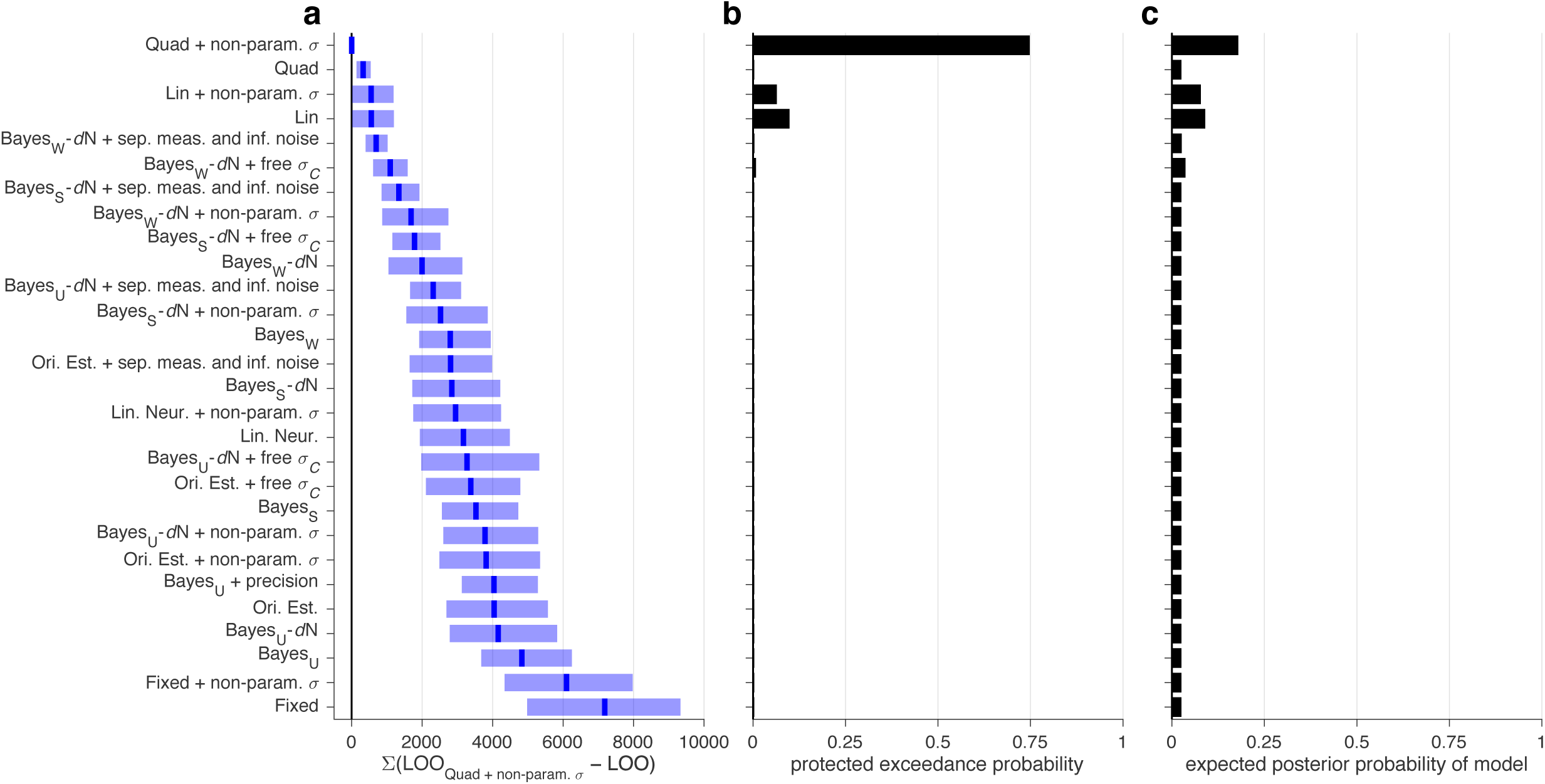
Model comparison, experiment 1. Models were fit jointly to Task A and B category and confidence responses. (**a**) Medians and 95% CI of bootstrapped sums of LOO differences, relative to the best model. Higher values indicate worse fits. (**b**) The protected exceedance probability, i.e., the posterior probability that a model occurs more frequently than the others^66^. (**c**) The expected posterior probability that a model generated the data of a randomly chosen subject^70^.

**Table S1:**
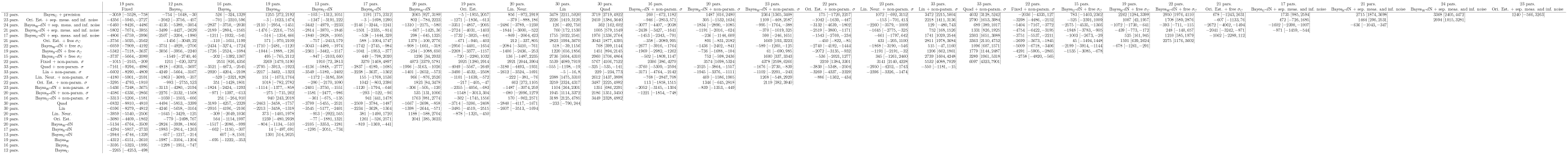
Cross comparison of all models in Figure S6. Cells indicate medians and 95% CI of bootstrapped summed LOO score differences. A negative median indicates that the model in the corresponding row had a higher score (better fit) than the model in the corresponding column. For readability, see instead *table_S1.xls*. For the parameters of each model, see *model_parameters.xls*.

**Figure S7:**
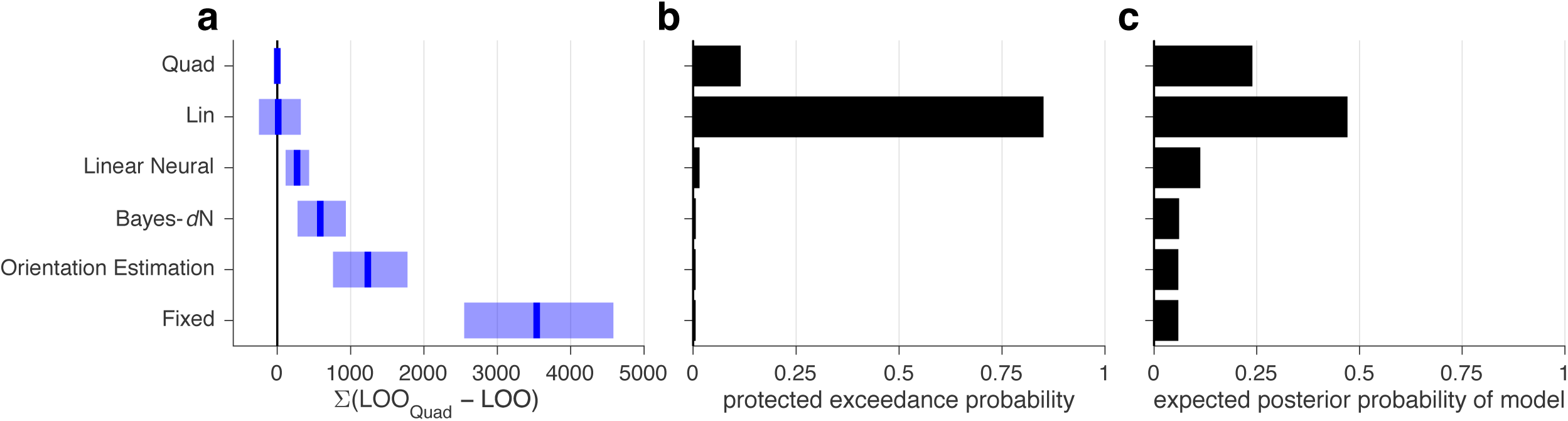
Model comparison, experiment 1. Models were fit to Task A category and confidence responses. See Figure S6 caption.

**Table S2:** Cross comparison of all models in Figure S7. See Table S1 caption.

|  |  | 12 pars.<br>Fixed | 13 pars.<br>Bayes-*dN* | 12 pars.<br>Ori. Est. | 13 pars.<br>Lin. Neur. | 16 pars.<br>Lin |
| --- | --- | --- | --- | --- | --- | --- |
| 16 pars. | Quad | -3534 [-4552, -2529] | -581 [-938, -278] | -1241 [-1798, -767] | -270 [-436, -117] | -14 [-325, 246] |
| 16 pars. | Lin | -3532 [-4353, -2651] | -572 [-799, -339] | -1232 [-1566, -863] | -259 [-609, 124] |  |
| 13 pars. | Lin. Neur. | -3255 [-4343, -2231] | -313 [-724, 75] | -972 [-1599, -412] |  |  |
| 12 pars. | Ori. Est. | -2302 [-2881, -1705] | 651 [425, 885] |  |  |  |
| 13 pars. | Bayes-*dN* | -2956 [-3723, -2163] |  |  |  |  |

**Figure S8:**
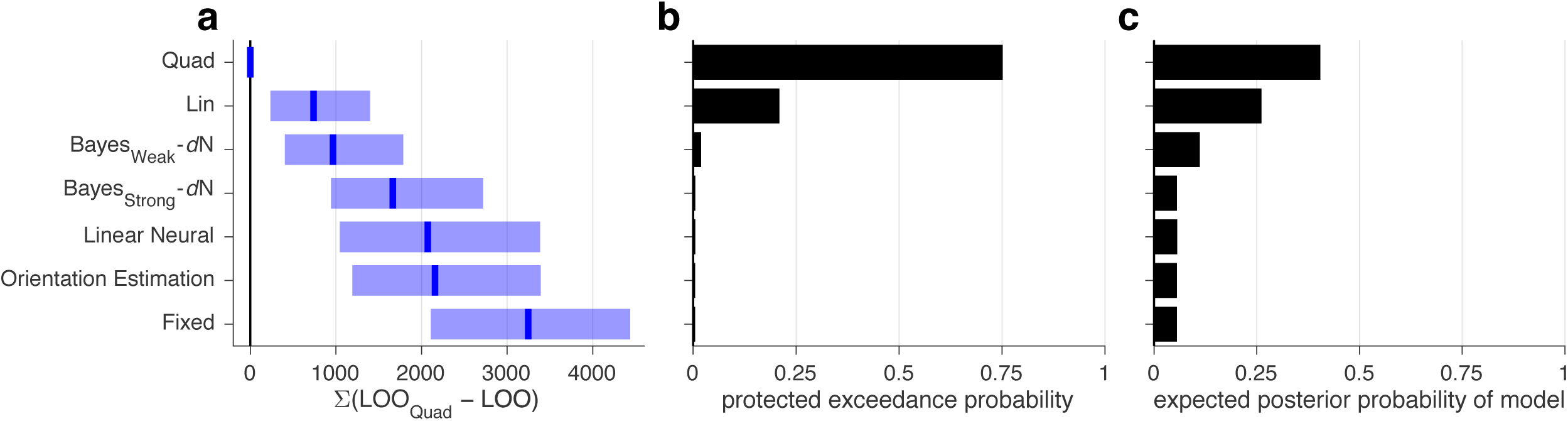
Model comparison, experiment 1. Models were fit to Task B category and confidence responses. See Figure S6 caption.

**Table S3:**
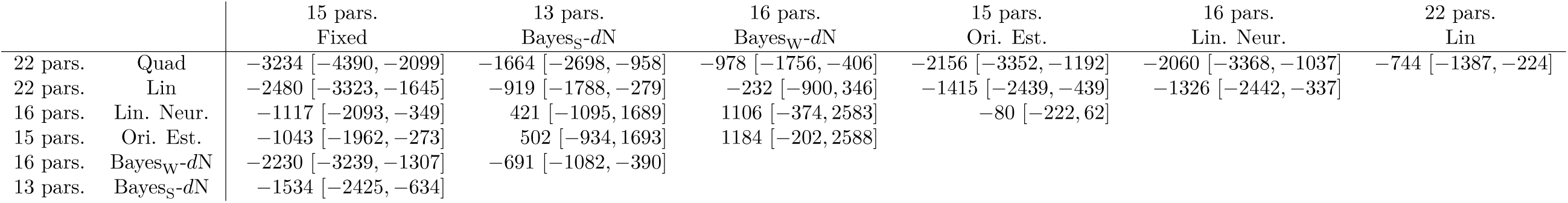
Cross comparison of all models in Figure S8. See Table S1 caption.

**Figure S9:**
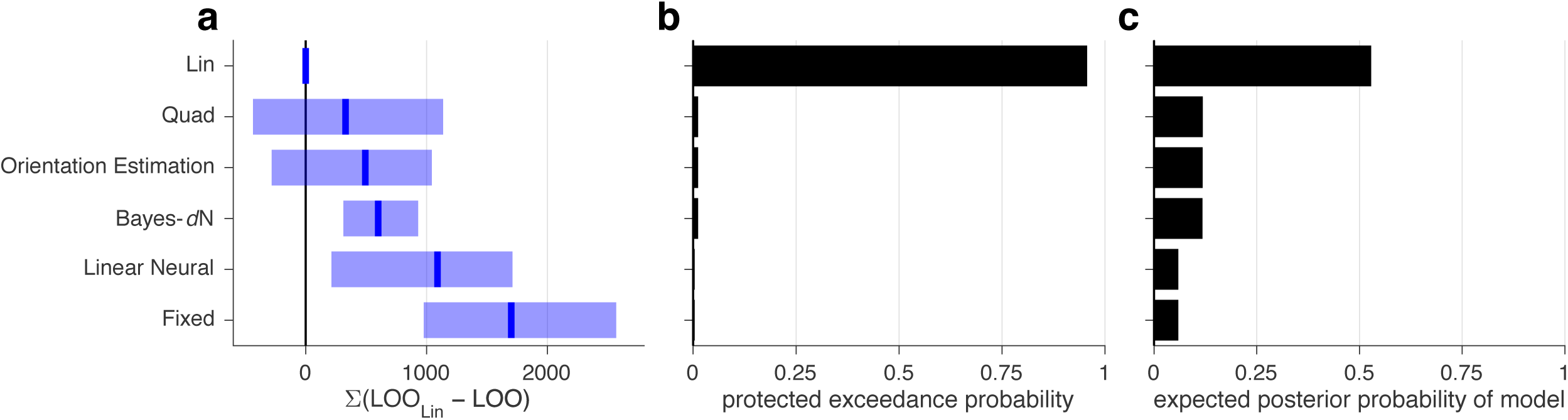
Model comparison, experiment 1. Models were fit jointly to Task A and B category choices. See Figure S6 caption.

**Table S4:**
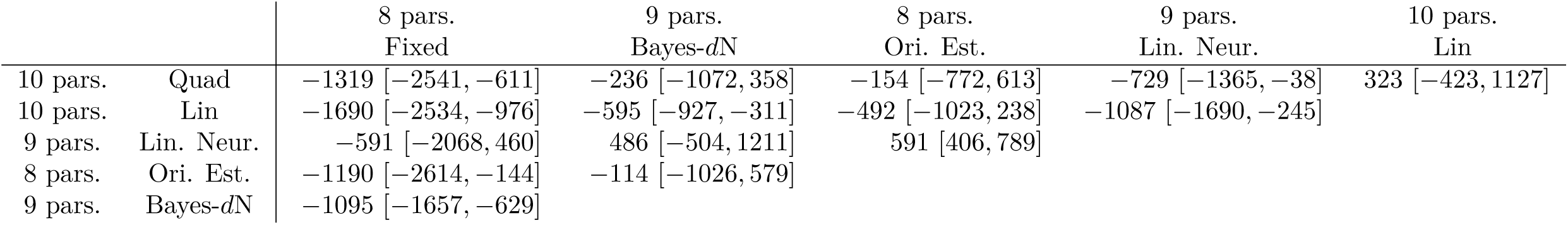
Cross comparison of all models in Figure S9. See Table S1 caption.

**Figure S10:**
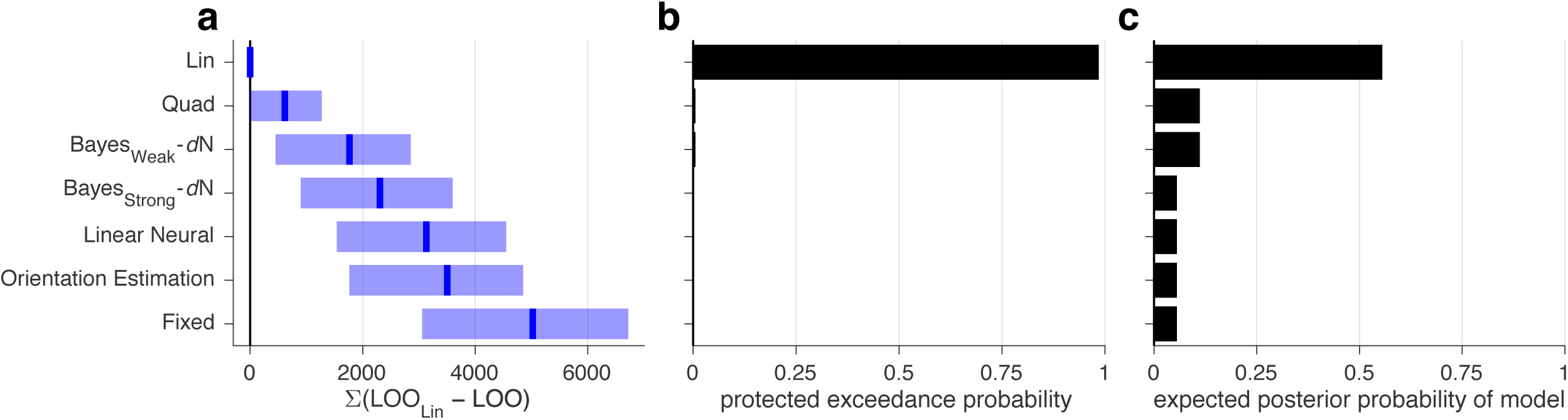
Model comparison, experiment 1. Noise parameters were fit to Task A category choices and then fixed during the fitting of Task B category and confidence responses. See Figure S6 caption.

**Table S5:**
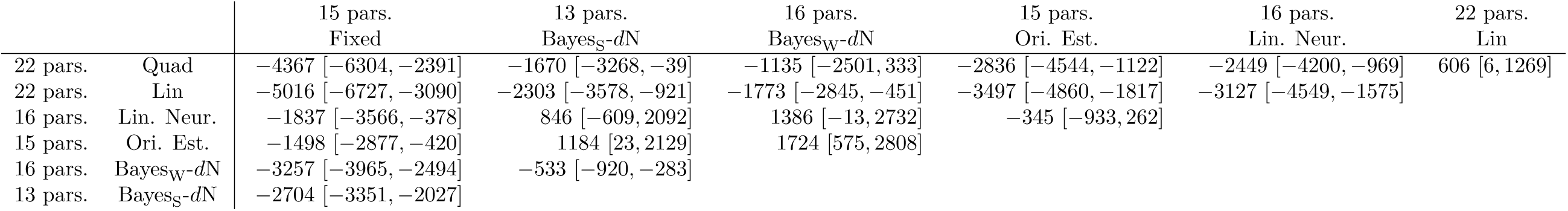
Cross comparison of all models in Figure S10. See Table S1 caption.

|  |  | 15 pars.<br>Fixed | 13 pars.<br>Bayes<sub>S</sub>-dN | 16 pars.<br>Bayes<sub>W</sub>-dN | 15 pars.<br>Ori. Est. | 16 pars.<br>Lin. Neur. | 22 pars.<br>Lin |
| --- | --- | --- | --- | --- | --- | --- | --- |
| 22 pars. | Quad | -4367 [-6304, -2391] | -1670 [-3268, -39] | -1135 [-2501, 333] | -2836 [-4544, -1122] | -2449 [-4200, -969] | 606 [6, 1269] |
| 22 pars. | Lin | -5016 [-6727, -3090] | -2303 [-3578, -921] | -1773 [-2845, -451] | -3497 [-4860, -1817] | -3127 [-4549, -1575] |  |
| 16 pars. | Lin. Neur. | -1837 [-3566, -378] | 846 [-609, 2092] | 1386 [-13, 2732] | -345 [-933, 262] |  |  |
| 15 pars. | Ori. Est. | -1498 [-2877, -420] | 1184 [23, 2129] | 1724 [575, 2808] |  |  |  |
| 16 pars. | Bayes<sub>W</sub>-dN | -3257 [-3965, -2494] | -533 [-920, -283] |  |  |  |  |
| 13 pars. | Bayes<sub>S</sub>-dN | -2704 [-3351, -2027] |  |  |  |  |  |

**Figure S11:**
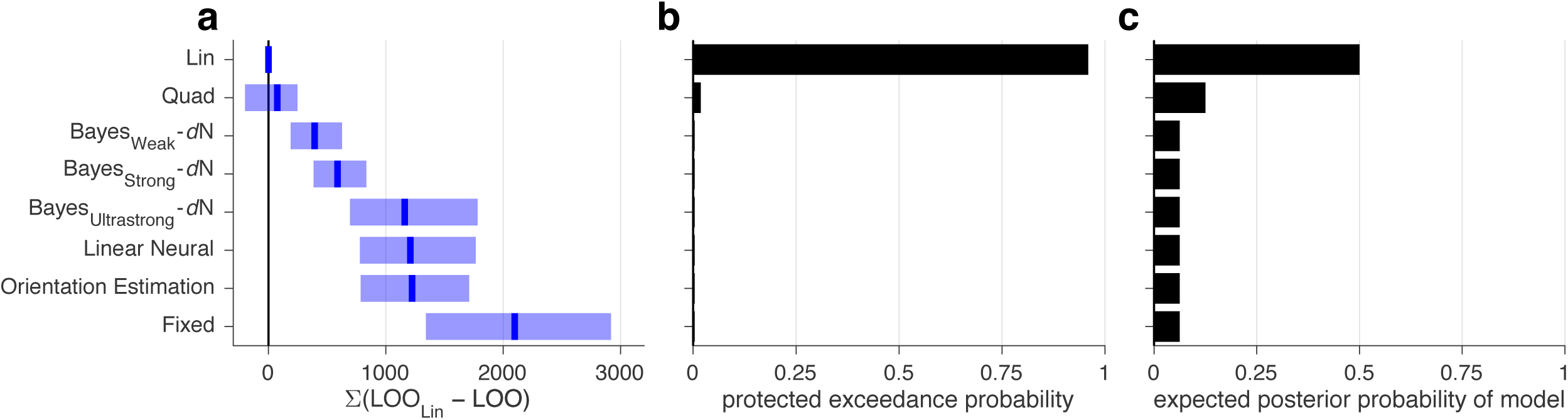
Model comparison, experiment 2. Models were fit jointly to Task A and B category and confidence responses. See Figure S6 caption.

**Table S6:** Cross comparison of all models in Figure S11. See Table S1 caption.

|  |  | 19 pars.<br>Fixed | 13 pars.<br>Bayes<sub>U</sub>-dN | 17 pars.<br>Bayes<sub>S</sub>-dN | 20 pars.<br>Bayes<sub>W</sub>-dN | 19 pars.<br>Ori. Est. | 20 pars.<br>Lin. Neur. | 30 pars.<br>Lin |
| --- | --- | --- | --- | --- | --- | --- | --- | --- |
| 30 pars. | Quad | -2014 [-3036, -1186] | -1096 [-1807, -530] | -523 [-893, -220] | -331 [-562, -109] | -1136 [-1815, -638] | -1124 [-1922, -613] | 74 [-195, 252] |
| 30 pars. | Lin | -2095 [-2889, -1344] | -1160 [-1780, -694] | -589 [-841, -375] | -396 [-622, -186] | -1218 [-1680, -791] | -1205 [-1757, -785] |  |
| 20 pars. | Lin. Neur. | -876 [-1401, -395] | 55 [-711, 693] | 623 [99, 1184] | 801 [253, 1491] | -7 [-219, 216] |  |  |
| 19 pars. | Ori. Est. | -872 [-1297, -487] | 56 [-561, 574] | 620 [218, 1089] | 813 [356, 1394] |  |  |  |
| 20 pars. | Bayes<sub>W</sub>-dN | -1678 [-2542, -1007] | -767 [-1262, -386] | -190 [-363, -82] |  |  |  |  |
| 17 pars. | Bayes<sub>S</sub>-dN | -1490 [-2210, -886] | -565 [-1032, -266] |  |  |  |  |  |
| 13 pars. | Bayes<sub>U</sub>-dN | -907 [-1572, -365] |  |  |  |  |  |  |

**Figure S12:**
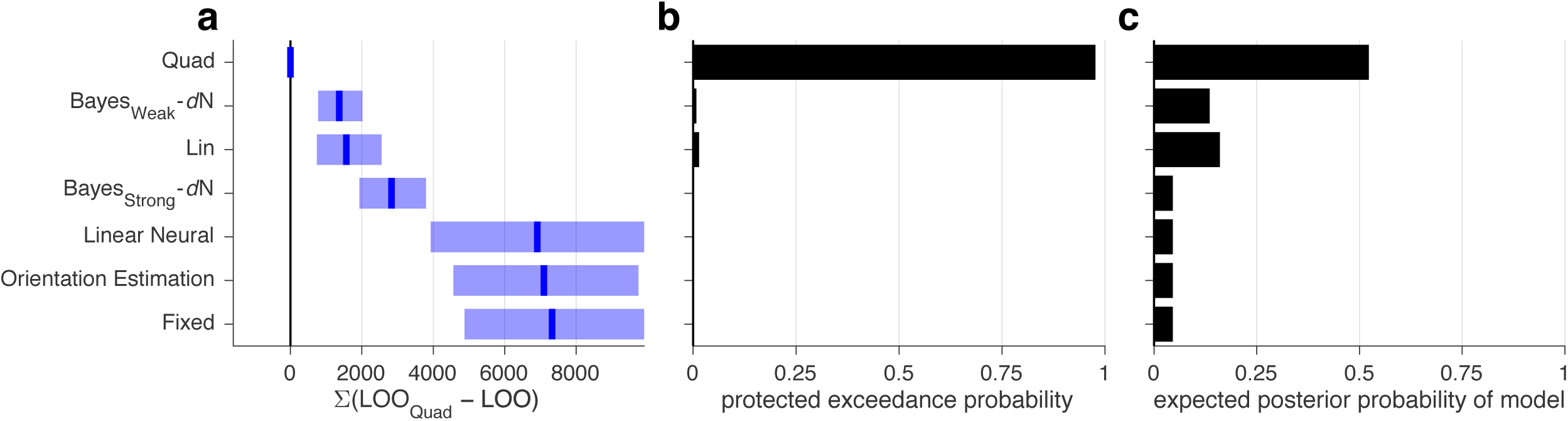
Model comparison, experiment 3. Models were fit to Task B category and confidence responses. See Figure S6 caption.

**Table S7:** Cross comparison of all models in Figure S12. See Table S1 caption.

|  |  | 15 pars.<br>Fixed | 13 pars.<br>Bayes<sub>S</sub>-dN | 16 pars.<br>Bayes<sub>W</sub>-dN | 15 pars.<br>Ori. Est. | 16 pars.<br>Lin. Neur. | 22 pars.<br>Lin |
| --- | --- | --- | --- | --- | --- | --- | --- |
| 22 pars. | Quad | -7326 [-9955, -4905] | -2833 [-3807, -1926] | -1361 [-2022, -777] | -7120 [-9838, -4636] | -6902 [-10376, -3981] | -1577 [-2562, -750] |
| 22 pars. | Lin | -5759 [-7866, -3694] | -1240 [-2567, 65] | 226 [-812, 1246] | -5530 [-7707, -3539] | -5337 [-8191, -2846] |  |
| 16 pars. | Lin. Neur. | -450 [-1535, 1290] | 4114 [733, 7796] | 5552 [2338, 9135] | -214 [-1176, 1256] |  |  |
| 15 pars. | Ori. Est. | -256 [-841, 423] | 4311 [1432, 7134] | 5727 [3067, 8527] |  |  |  |
| 16 pars. | Bayes<sub>W</sub>-dN | -5967 [-8702, -3369] | -1454 [-2179, -835] |  |  |  |  |
| 13 pars. | Bayes<sub>S</sub>-dN | -4505 [-7282, -1816] |  |  |  |  |  |

**Figure S13:**
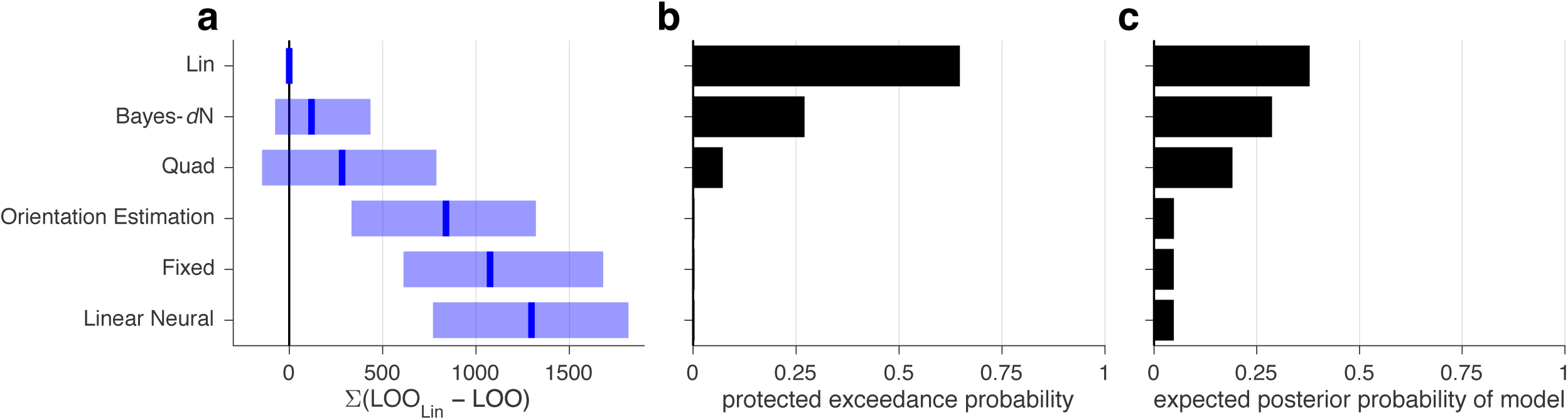
Model comparison, experiment 3. Models were fit to Task B category choices. See Figure S6 caption.

**Table S8:** Cross comparison of all models in Figure S13. See Table S1 caption.

|  |  | 7 pars.<br>Fixed | 8 pars.<br>Bayes-*dN* | 7 pars.<br>Ori. Est. | 8 pars.<br>Lin. Neur. | 8 pars.<br>Lin |
| --- | --- | --- | --- | --- | --- | --- |
| 8 pars. | Quad | -777 [-1361, -359] | 162 [-290, 670] | -531 [-1059, -23] | -988 [-1526, -549] | 290 [-138, 793] |
| 8 pars. | Lin | -1084 [-1675, -619] | -117 [-436, 76] | -830 [-1317, -334] | -1294 [-1825, -778] |  |
| 8 pars. | Lin. Neur. | 215 [-566, 827] | 1174 [535, 1772] | 457 [254, 685] |  |  |
| 7 pars. | Ori. Est. | -255 [-987, 369] | 707 [119, 1259] |  |  |  |
| 8 pars. | Bayes-*dN* | -964 [-1290, -663] |  |  |  |  |

**Figure S14:**
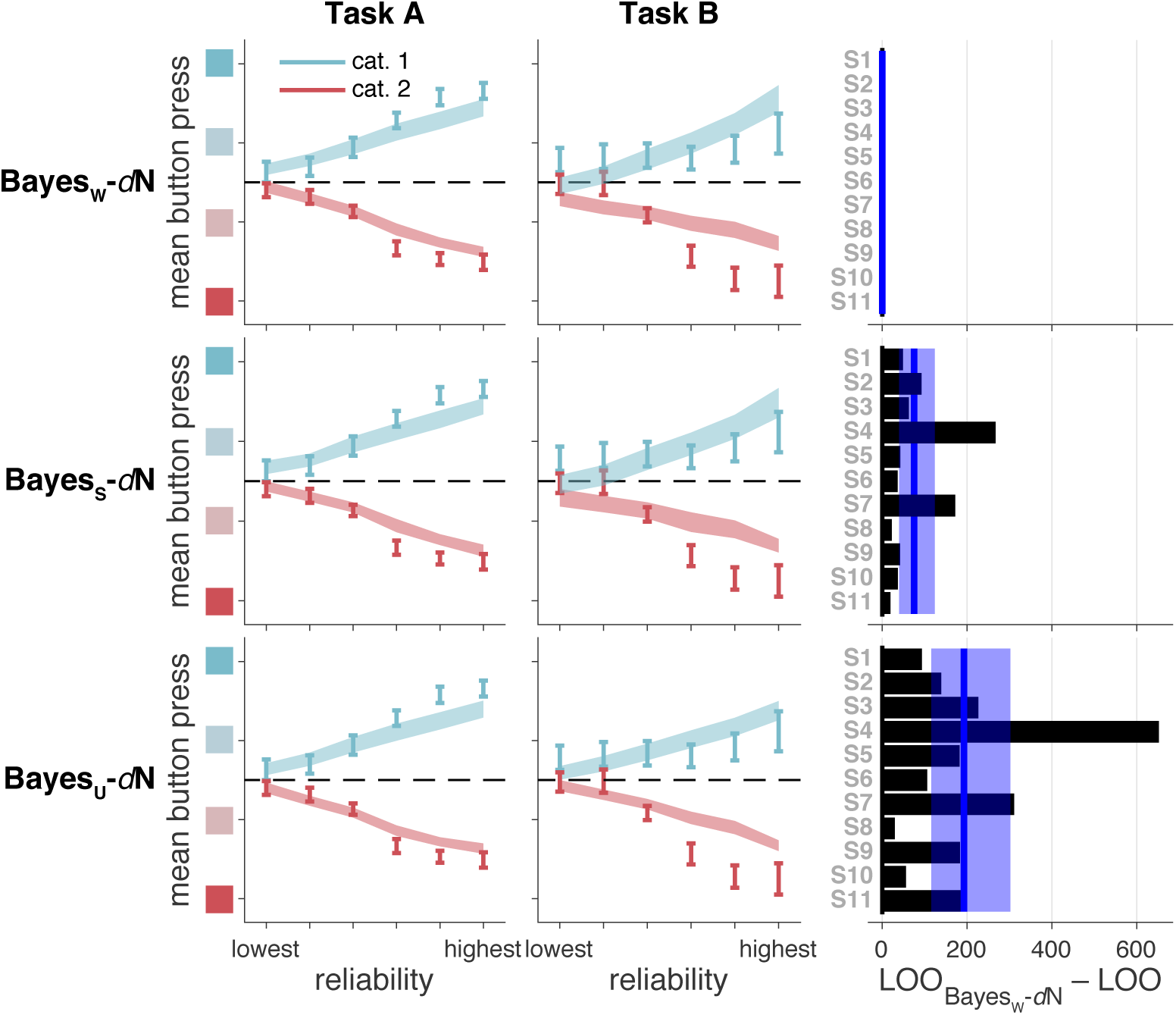
Model fits and model comparison for the three strengths of the Bayesian model, as in Figure 5. In the main text, Bayes_Weak_-*d*N is referred to simply as Bayes.

**Figure S15:**
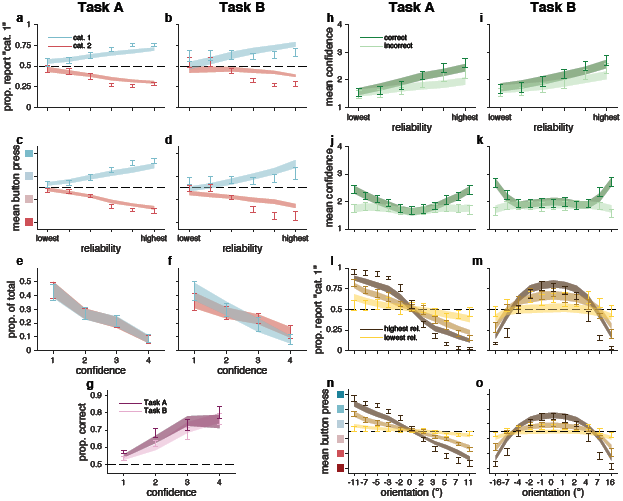
Bayes_Weak_-*d*N fits, as in Figure 3. In the main text, Bayes_Weak_-*d*N is referred to simply as Bayes.

**Figure S16:**
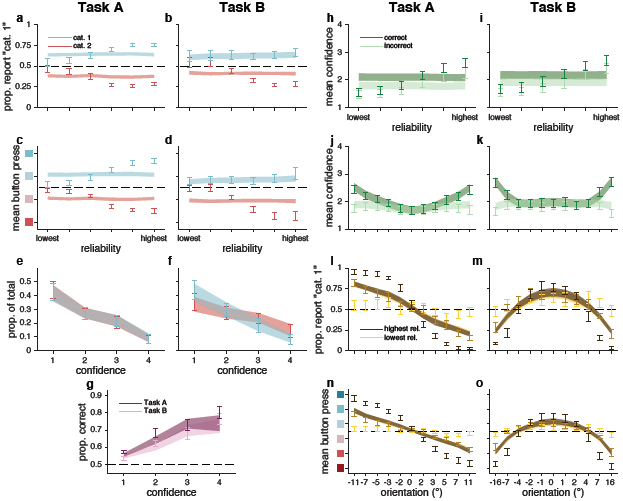
Fixed fits, as in Figure 3.

**Figure S17:**
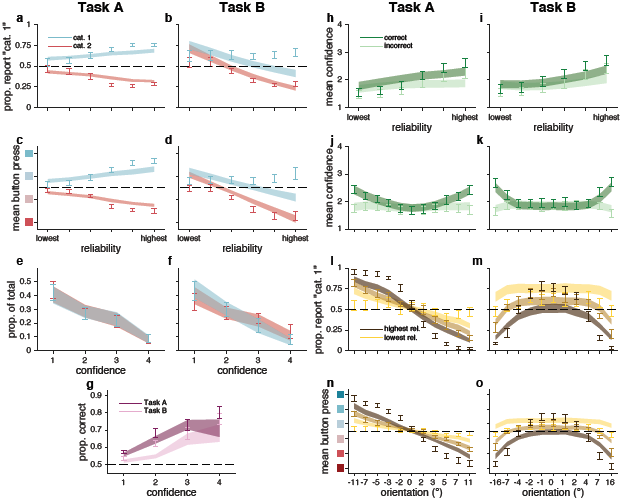
Orientation Estimation fits, as in Figure 3.

**Figure S18:**
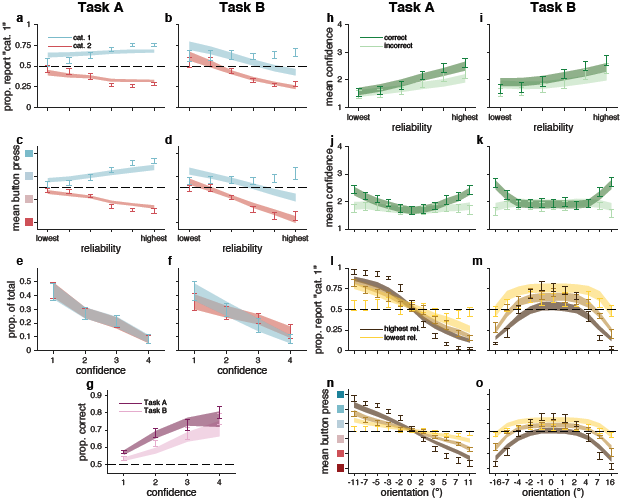
Linear Neural fits, as in Figure 3.

**Figure S19:**
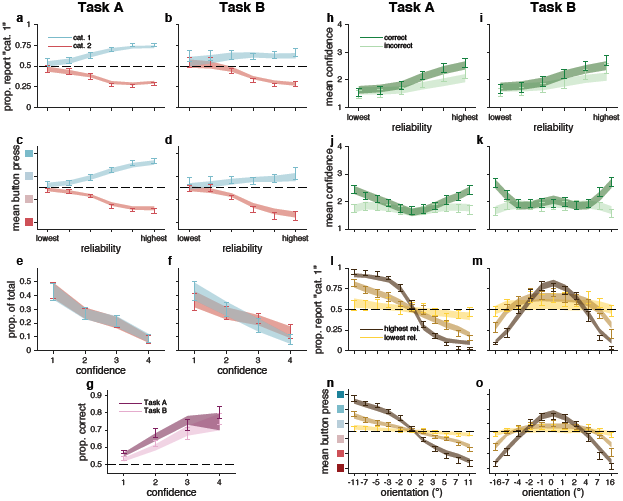
Lin fits, as in Figure 3.

